# Trunk sensory and motor cortex is preferentially integrated with hindlimb sensory information that supports trunk stabilization

**DOI:** 10.1101/2020.08.31.272583

**Authors:** Bharadwaj Nandakumar, Gary H. Blumenthal, Francois Philippe Pauzin, Karen A. Moxon

**Affiliations:** Department of Biomedical Engineering, Science, and Health Systems, Drexel University, Philadelphia, PA; Department of Biomedical Engineering, University of California- Davis, Davis, CA; Center for Neuroscience, Davis, CA

**Author notes:** these two authors contributed equally (Bharadwaj Nandakumar, Gary H. Blumenthal). Corresponding Author: Karen A. Moxon, PhD, University of California, Davis, 451 E. Health Sciences Drive, GBSF 3321, Davis, CA 95616.

**Keywords:** dermatome, dorsal root ganglion, motor cortex, mapping, sensory cortex

## Abstract

Sensorimotor integration in the trunk system has been poorly studied despite its importance for examining functional recovery after neurological injury or disease. Here, we mapped the relationship between thoracic dorsal root ganglia and trunk sensory cortex (S1) to create a detailed map of the extent and internal organization of trunk primary sensory cortex, and trunk primary motor cortex (M1) and showed that both cortices are somatotopically complex structures that are larger than previously described. Surprisingly, projections from trunk S1 to trunk M1 were not anatomically organized. We found relatively weak sensorimotor integration between trunk M1 and S1 and between trunk M1 and forelimb S1 compared to extensive integration between trunk M1 and hindlimb S1 and M1. This strong trunk/hindlimb connection was identified for high intensity stimuli that activated proprioceptive pathways. To assess the implication of this integration, the responses in sensorimotor cortex were examined during a postural control task and supported sensorimotor integration between hindlimb sensory and lower trunk motor cortex. Together, these data suggest that trunk M1 is guided predominately by hindlimb proprioceptive information that reached the cortex directly via the thalamus. This unique sensorimotor integration suggests an essential role for the trunk system in postural control, and its consideration could be important for understanding studies regarding recovery of function after spinal cord injury.

**Significance:** This work identifies extensive sensorimotor integration between trunk and hindlimb cortices, demonstrating that sensorimotor integration is an operational mode of the trunk cortex in intact animals. The functional role of this integration was demonstrated for postural control when the animal was subjected to lateral tilts. Furthermore, these results provide insight into cortical reorganization after spinal cord injury making clear that sensorimotor integration after SCI is an attempt to restore sensorimotor integration that existed in the intact system. These results could be used to tailor rehabilitative strategies to optimize sensorimotor integration for functional recovery.

## Introduction

Transmission of information between sensory and motor systems, or sensorimotor integration, is crucial for perception (1) and volitional control of movement (2). Understanding the substrates of sensorimotor integration is important for studies examining learning and recovery after neurological injury or disease. For example, sensorimotor integration has been extensively studied in the rodent whisker system(1, 3–8) giving rise to a better understanding of how rodents use their whiskers optimally to navigate and discriminate features of their environment. Furthermore, research on the forelimb (9–13) and hindlimb systems (14–18) have highlighted the importance of sensorimotor integration for appropriate locomotor function. These studies found extensive integration between anatomically and topographically corresponding sensory and motor cortices, with little cross-region integration (e.g. integration between whisker sensory and hindlimb motor cortices). Yet, little is known about sensorimotor integration within the trunk cortex or between the trunk cortex and other sensorimotor cortices.

Classic mapping studies of the rodent primary sensory cortex (S1) and primary motor cortex (M1) have roughly outlined the location and border of trunk S1 and M1 (14, 19, 20). More recently, studies added some distinctions within trunk S1, including a ventral trunk representation (21, 22) and a genital representation (23). Despite these findings, the internal somatotopy of trunk S1 remains ill defined, in part, due to the limited assessment of spinal dermatomes of the thoracic regions (24, 25). Similarly, the trunk M1 is mentioned in most mapping studies (26–28) and some information has emerged from recent studies examining cortical reorganization after spinal cord injury (29–35). However, little is known about the internal somatotopy of trunk M1 (30, 33, 34, 36). Further study of the somatotopy of trunk S1 and M1, as well as how these cortices integrate information is needed to more fully understand the role of trunk cortex both in naive animals and after neurological injury or disease (30, 32, 34, 35).

Thus, the aims of the current study were to define the somatotopy of trunk S1 and trunk M1 and examine sensorimotor integration of trunk cortex. First, to examine the internal organization of trunk S1, electrophysiological mapping was done at the spinal level to identify thoracic dermatomes and their corresponding representation in S1. Intracortical microstimulation (ICMS) was used to examine the extent and internal organization of trunk M1. Then, sensorimotor integration was assessed by examining sensory evoked potentials across broad regions of sensorimotor cortex. Retrograde tracing was then done to understand the source of sensory integration in trunk M1. Finally, to understand the functional role of sensorimotor integration, we recorded single neuron activity from trunk M1 and S1 in response to unexpected postural perturbations, while animals stood on a tilting platform. Results from mapping studies reveal an important somatotopic organization within both the trunk S1 and M1cortices. Furthermore, there is extensive sensorimotor integration between trunk and hindlimb systems compared to the relatively weak integration within trunk, and between trunk and forelimb cortices. Evidence from response latency and tracing studies suggest that this trunk/hindlimb sensorimotor integration is mediated by thalamo-cortical projections carrying proprioceptive information. Importantly, this integration of hindlimb proprioceptive information and trunk motor cortex is activated during postural adjustments to allow the animal to stabilize the trunk and maintain balance. These insights into trunk sensorimotor organization are important for understanding how information is processed during locomotion, as well as for efficiently adapting rehabilitative strategies after spinal cord injury.

## Results

### Dermatomes of upper thoracic DRGs overlap more than those of lower thoracic DRGs

To study how trunk sensory information is represented in the brain, it is important to understand how this sensory information is first represented at the spinal level. While the upper and lower thoracic dermatomes were previously mapped (24, 25, 37, 38), the mid thoracic dermatomes have not been mapped extensively in the rat, nor is the representation of these dermatomes in the cortex known. To this end, we recorded single neuron activity from the DRGs in the thoracic level (T1-T13) and mapped the thoracic dermatomes (Figure 1A). An average of 6 +/− 3 DRGs were recorded per animal (*n* =15) across the thoracic level. The thoracic dermatomes were rectangular bands with overlapping receptive fields that extended from the dorsal midline to the midline on the ventral side of the trunk. The T1-T3 dermatomes had receptive fields that extended into the forelimb, while the receptive fields of the remaining thoracic dermatomes were limited to the trunk (Figure 1B). The width of the thoracic dermatomes remained constant in the rostrocaudal direction along the body (one-way repeated measures ANOVA, *F* (12, 59) = 1.41, *p* = .44; Figure 1C), consistent with studies performed on cats (39, 40), sheeps (41), and monkeys (42, 43). However, the amount of overlap between adjacent dermatomes decreased significantly from rostral to caudal (one-way repeated measures ANOVA, *F* (11, 44) = 2.52, *p* < .05: Figure 1D). This decrease in overlap was due to a shift in the average center position of adjacent dermatomes (one-way ANOVA, *F* (2, 61) = 7.73, *p* < .01; Figure 1E). Tukey post hoc revealed a significant increase in average distance between the dermatomes of upper (T1-T5) and those of both the mid (T5-T9; *p* < .05) and the lower thoracic (T9-T13; *p* < .001) regions. Therefore, rostral DRGs appeared to integrate more information from neighboring dermatomes than caudal DRGs.

**Figure 1.**
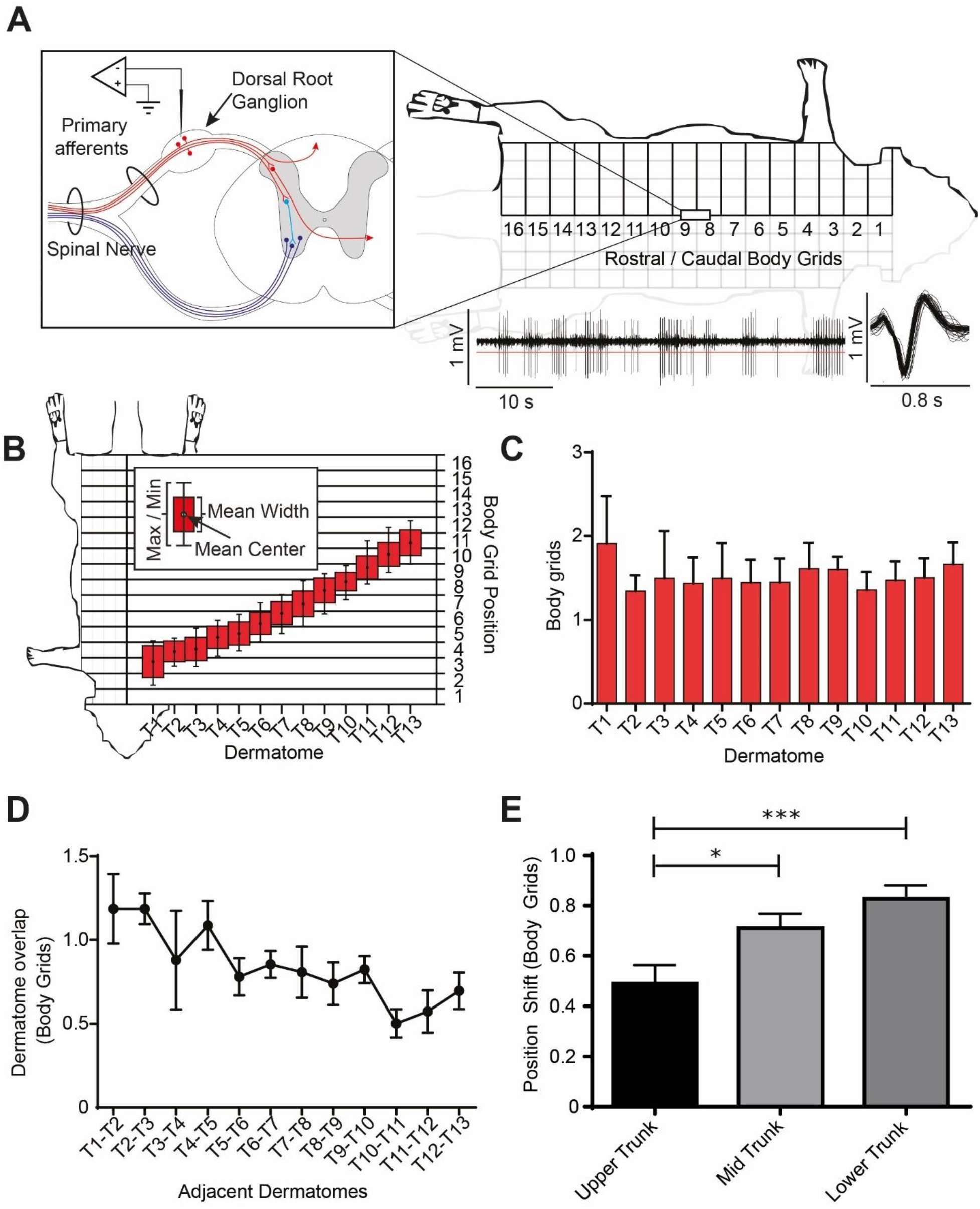
Trunk spinal dermatomes. A) Dermatome map methodological diagram. A tungsten microelectrode was inserted into the dorsal root ganglions (DRGs) to record from primary afferent cell bodies and receptive fields were determined. An example of a continuous neural trace and a sorted single unit waveform are shown in the bottom right part. B) Average dermatome width (in body grid units) and center position plotted along the rostrocaudal axis of the body. The error bar represents the most rostral and the most caudal body grid positions of each dermatome across all animals. C) Average dermatome width is similar throughout the rostrocaudal axis. D) Average overlap in body grids unit between adjacent dermatomes showed a shift in the rostrocaudal axis. E) Average distance in between neighboring dermatomes within the upper (T1-T5), mid (T5-T9), and lower (T9-T13) thoracic dermatomes showed a shift in the rostrocaudal axis, with a significant difference for the average distance in between neighboring dermatomes between upper trunk and mid trunk and between upper trunk and lower trunk.

### Information from mid and lower trunk are well integrated in trunk S1

A somatotopic map of the trunk and surrounding sensory cortices was constructed using the information from the dermatomes. In each cortical location, the proportion of cells responding to each body category was calculated (Figure. 2A). An average of 9 +/− 3 cortical locations were sampled per animal (N=40 animals), with an average of 8 +/− 3 single neurons sampled per location. In total, more than 2900 neurons were recorded. The trunk sensory cortex was located along the caudal edge of forelimb and hindlimb S1 consistent with previous studies in rats (19, 21, 22). The representation of the neck was most lateral, with the tail representation most medial (Figure 2B). Dorsal trunk S1 was located more caudal to ventral trunk S1. The ventral trunk S1, consistent with previous studies (21, 22, 44), was nestled between the forelimb S1 (lateral) and the hindlimb S1 (medial), and rostral to midthoracic (T6-T9) trunk representations, overlapping with the genital cortex described in previous studies (23). This extensive mapping suggests that the trunk representation is larger than previously reported (14, 26, 29, 34).

**Figure 2.**
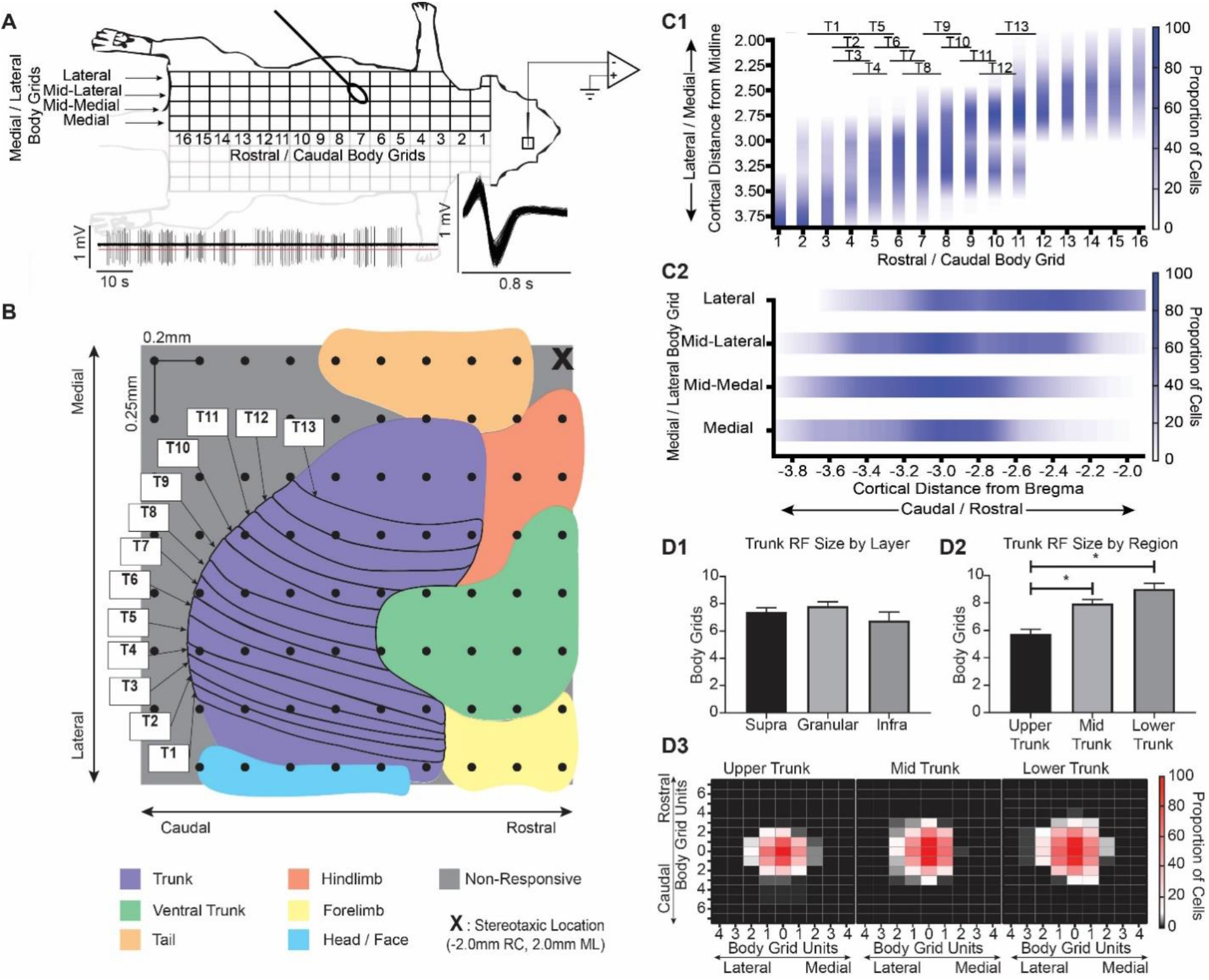
Relationship of Trunk S1 organization in relationship to spinal dermatomes. A) Sensory map methodological diagram. A tungsten microelectrode was inserted into several locations within and around the trunk sensory cortex. Single units were isolated, and their receptive fields were determined. Example of a continuous neural trace and a sorted single unit waveform can be seen at the bottom part. B) Cortical representation of the thoracic dermatomes. The map shows the average cortical representations surface for all cortical layers. 2920 neurons were recorded to construct the map. C1) Proportion of cells identified in the medio-lateral cortical axis across all animals, associated with body grid rows in which the cortical cells are responsive to light tactile stimulation. A higher proportion of rostral trunk RFs were responsive in lateral cortical coordinates while a higher proportion of caudal trunk RFs were responsive in medial cortical coordinates. The rostrocaudal extent of the thoracic dermatomes relative to the body grid rows are also displayed. C2) Proportion of cells identified in the rostrocaudal axis across all animals, associated with body grid columns in which the cortical cells are responsive to light tactile stimulation. A higher proportion of lateral trunk RFs were responsive in rostral cortical coordinates while a higher proportion of medial trunk RFs were responsive in caudal cortical coordinates. D1) Trunk receptive field size (body grids units) for the supragranular, granular and infragranular layers are of similar size,. D2) Receptive field size is significantly different for the upper, mid and lower trunk S1 regions. D3) Receptive field centers are normalized to position (0, 0) and the proportion of cells responsive to the surrounding body grids are calculated and showed significant differences in size (refer D2).

Within the trunk representation, the thoracic dermatomes were represented from T1, laterally, to T13, medially, consistent with a study in humans (45). As might be expected, there was extensive overlap of the cortical representation of neighboring thoracic dermatomes (Figure 2C). The rostrocaudal dimension of the dorsal trunk body was represented along the mediolateral axis of the cortex, with rostral trunk body represented laterally in the trunk S1 (Figure 2C1). The mediolateral dimension of the dorsal trunk body was represented in the rostrocaudal axis, with the most lateral part of dorsal trunk body represented rostrally in the cortex, just caudal to the ventral trunk representation (Figure 2C2). Unlike other sensory systems such as whisker and limbs that tend to have RF size differences across layers (11), the RF size of neurons in trunk S1 were similar across layers (one-way ANOVA, *F* (2, 479) = 1.45, *p* = .23; Figure 2D1). However, the RF size of trunk neurons did differ across the different regions of the trunk sensory cortex (one-way ANOVA, *F* (2, 434) = 19.71, *p* < .0001; Figure 2D2) with upper trunk neurons having smaller RF size compared to either mid or lower trunk neurons (5.7 +/− 3.7, 8.0 +/− 3.8, 9.0 +/− 5.0 body grids, respectively, Tukey post-hoc, p<0.001; Figure 2D3). This RF size analysis suggests that sensory information ascending from the thalamus is spread across large parts of trunk S1 early, immediately upon arrival in layer IV, with mid and lower trunk sensory information better integrated than that of upper trunk.

### Preferential integration of hindlimb and trunk sensory input within S1

To understand the integration of trunk, forelimb, and hindlimb sensory information, multichannel recordings were performed in the trunk, forelimb, and hindlimb S1 in response to peripheral electrical stimulation of the mid trunk (MT), forelimb (FL), and hindlimb (HL). To ensure fair comparison between the responses to the different stimulus locations on the body, the amplitudes of the sensory evoked potential (SEP) from each cortical region recorded from the granular layer to graded peripheral electric stimulation of each respective region (FL, HL and MT) were compared (Figure 3A). As expected, there was a significant increase in the SEP amplitude associated with increases in stimulus current regardless of stimulus location (two-way repeated measures ANOVA, *F* (3, 71) = 18.06, *p* < .01; Figure 3A). However, across stimulus location the SEP amplitudes were similar (two-way repeated measures ANOVA, *F* (2, 21) = 0.91, *p* = .41; Figure 3A), suggesting that the stimulus at each location activated the homologous cortical region similarly and that comparisons could be made between responses recorded from different brain regions to stimulation of the same location on the body.

**Figure 3.**
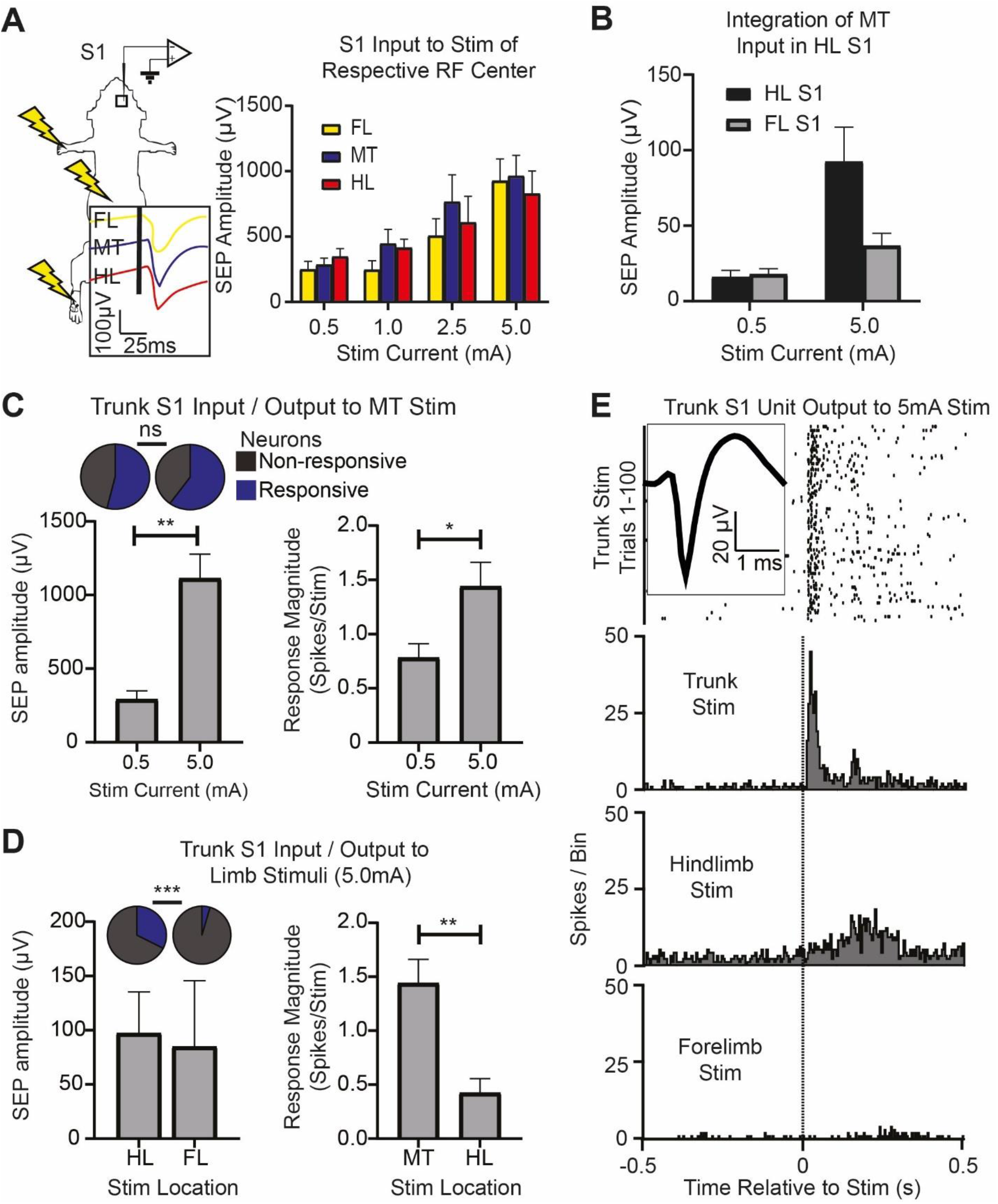
Sensory integration within trunk S1. A) Electric stimulation methodological diagram. Multichannel recordings were performed in the trunk, forelimb, and hindlimb S1 in response to peripheral electrical stimulation to the mid trunk (MT), forelimb (FL), and hindlimb (HL). The SEP from each cortical region recorded from the granular layer to graded peripheral electric stimulation of each respective region (FL, HL, MT) were compared. B) SEP amplitude in the forelimb S1 and hindlimb S1 in response to the low intensity (0.5 mA) and the high (5 mA) mid trunk stimulation. C) The relationship between sensory inputs (SEP amplitude in the granular layer, left, Mann-Whitney test, *p* < .01) and outputs (single neuron activity in all layers, right) in trunk S1 in response to low and high intensity MT stimuli. The inset on the top left represent the proportion of responsive cells for each stimuli. D) Bottom left: SEP amplitudes in trunk S1 in response to high intensity FL and HL stimulation. Top left: Proportion of trunk S1 neurons responding to hindlimb or forelimb stimulation. Right: Trunk S1 response to hindlimb and mid trunk stimulation (Mann-Whitney test, *p* < .01). E) Example PSTHs for HL, FL stimulation (5 mA) in trunk S1, illustrating that trunk S1 activity is modulated more by hindlimb than forelimb.

To understand the integration of trunk sensory information in S1, the amplitude of the SEP response to MT stimulation recorded from forelimb S1 was compared to the SEP response recorded from hindlimb S1. There were no significant differences in the SEP amplitude in response to low intensity trunk stimulation (0.5 mA, Mann-Whitney test; *p* = .79), however, the SEP amplitude to high intensity MT stimulation (5.0 mA) in hindlimb S1 was significantly greater than that recorded from forelimb S1 (Mann-Whitney test, *p* = .06; Figure 3B). Therefore, there is preferential integration of high intensity trunk sensory input in the hindlimb S1.

Next, within trunk S1, the relationship between inputs to layer IV cells (SEP amplitude) and outputs of trunk S1 (single neuron firing rate or proportion of responding neurons) in response to low and high intensity MT stimuli were examined to assess the effectiveness of the information transfer from input to output (Figure 3C). As noted above, SEP amplitude to high intensity MT stimulation was significantly greater than the response to low intensity MT stimulation (Mann-Whitney test, *p* < .01). This increase in input results in a greater magnitude of the response (spikes per stimulus) to high intensity stimuli (Mann-Whitney test, *p* < .05; Figure 3C) without a change in the proportion of responsive neurons (χ2 (1, 55) = 0.46; *p* = .50), suggesting the same cells are responding to low intensity stimuli, likely tactile, as those that respond to high intensity, likely proprioceptive.

Next, the contribution of high intensity FL and HL stimulation to the response in trunk S1 was examined. The SEP amplitude in trunk S1 to FL stimulation was similar to that of HL stimulation (Mann-Whitney test; p = .35). However, the proportion of neurons in trunk S1 that responded to HL stimulation was greater than the proportion responding to FL stimulation (χ2 (1, 84) = 11.16, p < .001; Figure 3D), suggesting that the transfer of incoming sensory information to output is more effective for HL stimulation than FL stimulation. In fact, too few cells responded to FL stimulation to allow any further analysis. As expected, the response of trunk S1 neurons to HL stimulation was smaller than the response to MT stimulation (Mann-Whitney test, *p* < .01; Figure. 3D, 3E), These results, taken together, suggest that extensive integration of high intensity sensory information between the hindlimb and trunk sensory cortices influences trunk S1 output. In the last section of the paper, we explore how this organization is used to encode the cortical response to unexpected tilts in the lateral plane.

### Trunk motor cortex is better integrated with hindlimb than forelimb motor cortex

To gain a better understanding of sensorimotor integration within the trunk cortex, it was essential to examine the extent and organization within trunk M1. Trunk M1 was mapped using intracortical microstimulation (ICMS) and movement representations were examined by analyzing movement and EMG responses from trunk and limb musculature (Figure 4A). Each of the 88 cortical locations were sampled on an average of 7 +/− 2 times, across 21 animals. Each animal contributed to the data with an average of 27 +/− 2 cortical locations per animal. The average threshold current was 51.3 +/− 23.4 μA. The areas of the cortex that most likely activated trunk musculature were within 1.5 mm lateral to midline. The rostrocaudal extent ranged from 0.25 mm rostral to bregma to 2.25 mm caudal to bregma (Figure 4B). This placed the rat trunk motor cortex medial to forelimb M1 and hindlimb M1 and just caudal to whisker M1.

**Figure 4.**
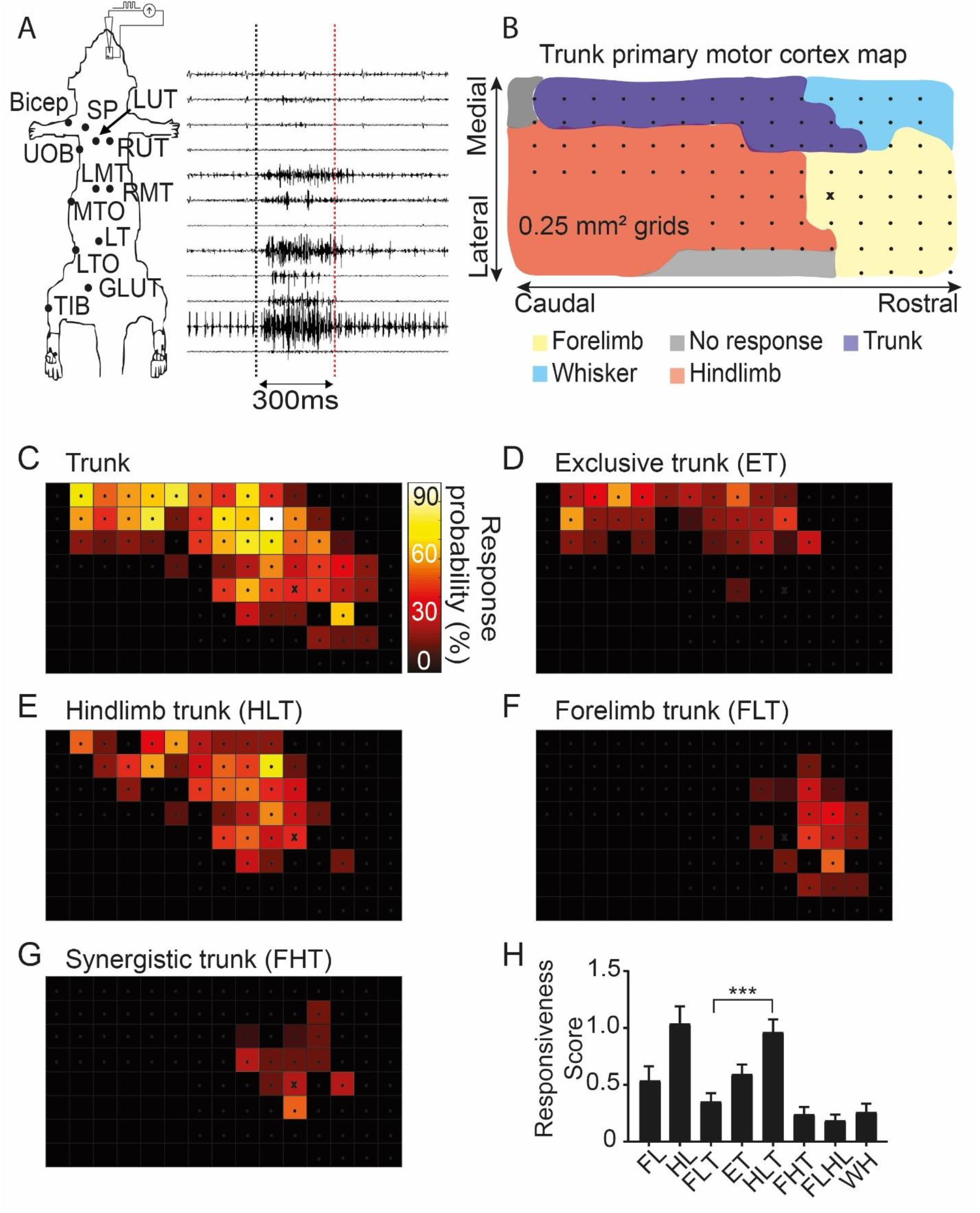
Coactivation of trunk musculature with forelimb and hindlimb. A) ICMS methodology for motor maps. Motor maps were obtained by intracortical current microsimulation (ICMS) in the infragranular layer of motor cortex. Evoked muscle activity is recorded through EMG electrodes implanted along the trunk, forelimb and hindlimb musculature (Top to bottom: Forelimb bicep (bicep), Spinous trapezius (SP), Left upper thoracic longissimus (LUT), Right upper thoracic longissimus (RUT), Upper external oblique (UOB), Left mid thoracic longissimus (LMT), Right mid thoracic longissimus (RMT), Mid thoracic external oblique (MTO), Lower thoracic longissimus (LT), Lower thoracic external oblique (LTO), Gluteus maximus (Glut), Tibialis anterior (Tib)). Observed movement and evoked muscle activity at threshold current is used to determine movement representation. B) Topography of trunk motor cortex based on most predominant response across animals. The dots refer to penetration locations sampled across animals. The X location refers to 0 mm RC, 2 mm ML relative to bregma. C) Proportion of penetrations that activates trunk musculature. D) Proportion of penetrations that activates trunk musculature exclusively. E) Proportion of penetrations that activates trunk and hindlimb musculature F) Proportion of penetrations that coactivates trunk and forelimb. G) Proportion of penetrations that coactivates trunk and both forelimb and hindlimb. H) Average responsiveness scores within trunk M1 is calculated for the different movement representation identified during mapping with ICMS.

A much larger area simultaneously activated trunk and other parts of the body, suggesting that trunk motor cortex is well integrated with forelimb and hindlimb motor cortex. In fact, it is possible to identify distinct coactivation zones between trunk and other parts of the body. The overall extent of the trunk coactivating with other parts of the body (Figure 4C) spanned −2.25 mm to 0.75 mm RC, and 1 mm to 2.5 mm ML relative to bregma, which is much larger than previously reported (26, 27, 33, 34, 36, 46). Interestingly, areas that exclusively activated trunk musculature (ET) within any given animal overlapped with trunk coactivation zones but were restricted to within 1.5 mm lateral to midline (Figure 4D). For each animal, the specific area of ET was quite small, and the position of ET relative to bregma was not consistent across animals. This suggests that there are likely to be few conditions under which trunk musculature is activated independently of the musculature of other parts of the body.

Despite this small area devoted to ET, coactivation of trunk with hindlimb musculature (HLT) was quite large (Figure 4E) and, not surprisingly, caudal to locations overlapping with forelimb (FLT, Figure 4F). In addition, consistent with an earlier study (47), in approximately half of the animals (45%), FL, HL and trunk (synergistic trunk or FHT) coactivated in locations between the HLT and FLT representation (Figure 4G). In order to quantify and compare the different movement representations found within the trunk coactivation zone, responsiveness score (34, 48) that represented the proportion of responses for each representation were compared. The responsiveness scores were different across coactivation zones (one-way ANOVA *F* (7,424) = 11.10, *p* < .001; Figure 4H). Importantly, the responsiveness score of HLT was significantly greater than FLT (Tukey’s multiple comparison test, *p* < .01) meaning that trunk coactivates more with hindlimbs across a larger region of cortex compared to forelimbs. Moreover, the responsiveness score of FLHL and FHT were very similar. These results demonstrated a topography within trunk motor cortex: a larger representation of trunk motor cortex compared to previous studies, and extensive integration of trunk motor output with the hindlimb motor cortex compared to the forelimb motor cortex.

### Trunk motor cortex is somatotopically organized

To understand the trunk musculature recruitment associated with the different coactivation zones, the EMG responses were examined in more detail. As expected, stimulation of forelimb trunk cortex (FLT) preferentially activated spinous trapezius (SP) and contralateral upper thoracic longissimus (LUT). FLT coactivation zone is thus responsible for upper thoracic trunk muscles activation (Figure 5A, 5B, 5C). Similarly, stimulation of hindlimb trunk cortex (HLT) activated the obliques along the mid and the lower thoracic level and therefore HLT coactivation zone is preferentially responsible for mid and lower trunk muscles activation (Figure 5A, 5B, 5C). Interestingly, stimulation of exclusive trunk (ET) cortex also activated the oblique but at all thoracic levels, suggesting that ET is important to coordinate movements of the entire trunk. Finally, stimulation of the synergistic trunk cortex (FHT) activated mostly trunk musculature at the mid thoracic level (Figure 5C). Therefore, the different trunk coactivation zones differentially activate segmental trunk muscles (upper, mid and lower thoracic levels) providing topography to trunk M1 motor control.

**Figure 5.**
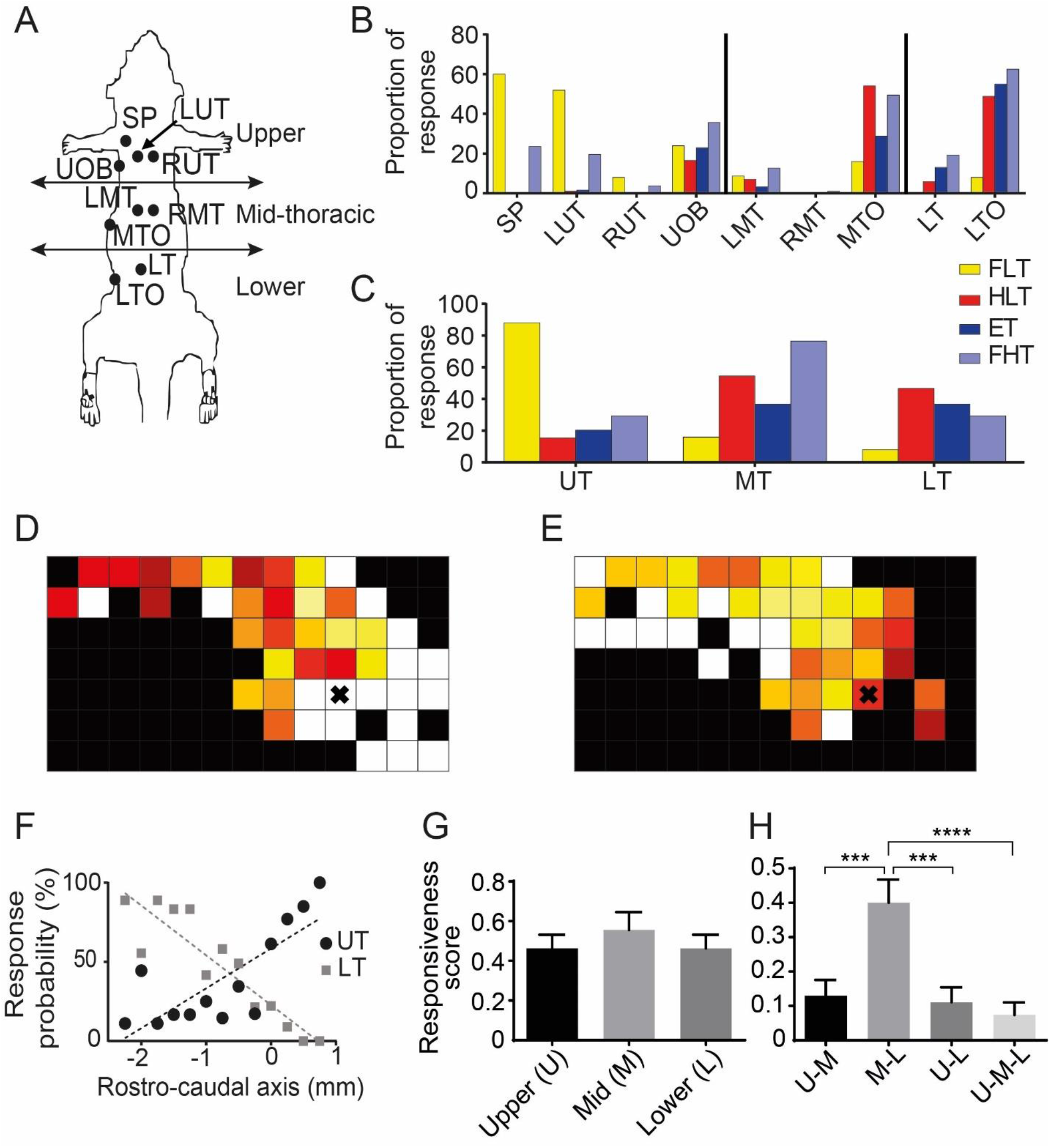
Recruitment of trunk musculature in the different coactivation zones. A) Methodological diagram showing EMG electrode locations of trunk muscles categorized into three groups along the rostrocaudal axis of the body: upper-thoracic, mid-thoracic and lower thoracic trunk muscles. B) Proportion of muscle responses in the different coactivation zones by muscle. C) Same graph as B but muscles are grouped within the three segmental zones seen in A. Upper-thoracic muscles are activated when FLT coactivation zone is stimulated and lower thoracic muscles are activated when HLT coactivation zone is stimulated. D) Proportion of penetrations that activated upper trunk musculature. E) Proportion of penetrations that activated lower trunk musculature. The X location refers to 0 mm RC, 2 mm ML relative to bregma. F) Graph showing the proportion of muscle responses based on visual observation & EMG responses of either upper or lower trunk musculature averaged across the rostrocaudal axis, lower thoracic G) Average responsiveness score (see method) in the trunk M1 for the different segmental zone: upper-, mid- and lower thoracic zones. H) Differences in the likelihood of segmental coactivation.

To gain more insight, we constructed two maps of trunk coactivation zone: the first to identify the proportion of penetrations across animals that activated upper trunk muscles (Figure 5D) and the second to identify the proportion that activated lower trunk muscles (Figure 5E). The mediocaudal region of trunk coactivation zone preferentially controlled lower thoracic trunk musculature while the rostrolateral region controlled upper thoracic trunk musculature. The lower trunk musculature was more influenced by the rostrolateral area of trunk coactivation zone and upper trunk musculature by the mediocaudal area of trunk coactivation zone. To demonstrate this topography along the rostrocaudal axis, the proportion of penetrations activating upper or lower thoracic trunk from mediolateral locations were averaged (Figure 5F). As one moves more rostral, there was an increase in the probability of activating upper trunk (*R^2^* = .61, *F* (1, 11) = 17.82, *p* < .01). Alternatively, moving caudally, there is an increase in the probability of activating lower trunk (*R^2^* = 0.83, *F* (1, 11) = 54.70, *p* < .01), demonstrating a clear somatotopy within the trunk coactivation zone.

Since segmental trunk muscles were differentially activated within this trunk coactivation zone, the amount and extent of activation within the coactivation zone was examined using the responsiveness score. The responsiveness scores were similar across the mid, upper and lower thoracic segmental levels (one-way ANOVA *F* (2,159) = 0.49, *p* = .54; Figure 5G), suggesting that the probability of cortex to activate the different segmental levels is similar. However, despite this similarity, there were differences in the likelihood of segmental coactivation (one-way ANOVA *F* (3,212) = 9.06, *p* < .001; Figure 5H) with the mid and lower thoracic muscles more likely to coactivate than other segmental muscle groups (Tukey’s multiple comparison test, *p* < .001). In summary, most of trunk cortex is devoted to activation with other regions of the body and cortical representation of mid and lower thoracic trunk muscles are integrated with hindlimb muscle representation while upper thoracic trunk muscles are integrated with forelimb muscle representation. These results were confirmed by synergy analysis using the amplitude of evoked EMG responses obtained from the different trunk musculature (Supplementary Figure 1).

### Sensory input to trunk motor cortex is dominated by hindlimb information

Given our understanding of integration of trunk sensory and motor cortex with cortical regions representing other parts of the body, we examined the integration of trunk M1 with sensory input from the limbs by recording the response in trunk M1 supragranular and infragranular layers in response to electric stimulation of FL, HL, MT, and UT (Figure 6A). There was little to no response in trunk M1 to light tactile, low intensity stimulation (0.5 mA) applied to any of the four body locations. However, this was not the case for high intensity stimulation (5.0 mA). Surprisingly, the SEP amplitude recorded from trunk motor cortex to high intensity sensory stimulation of the HL was greater than the SEP amplitude to stimulation of UT or MT (Fig. 6B, 6C). The responses to FL stimulation were similar to that of UT and MT stimulation, solidifying that trunk M1 is preferentially innervated with sensory information from hindlimbs.

**Figure 6.**
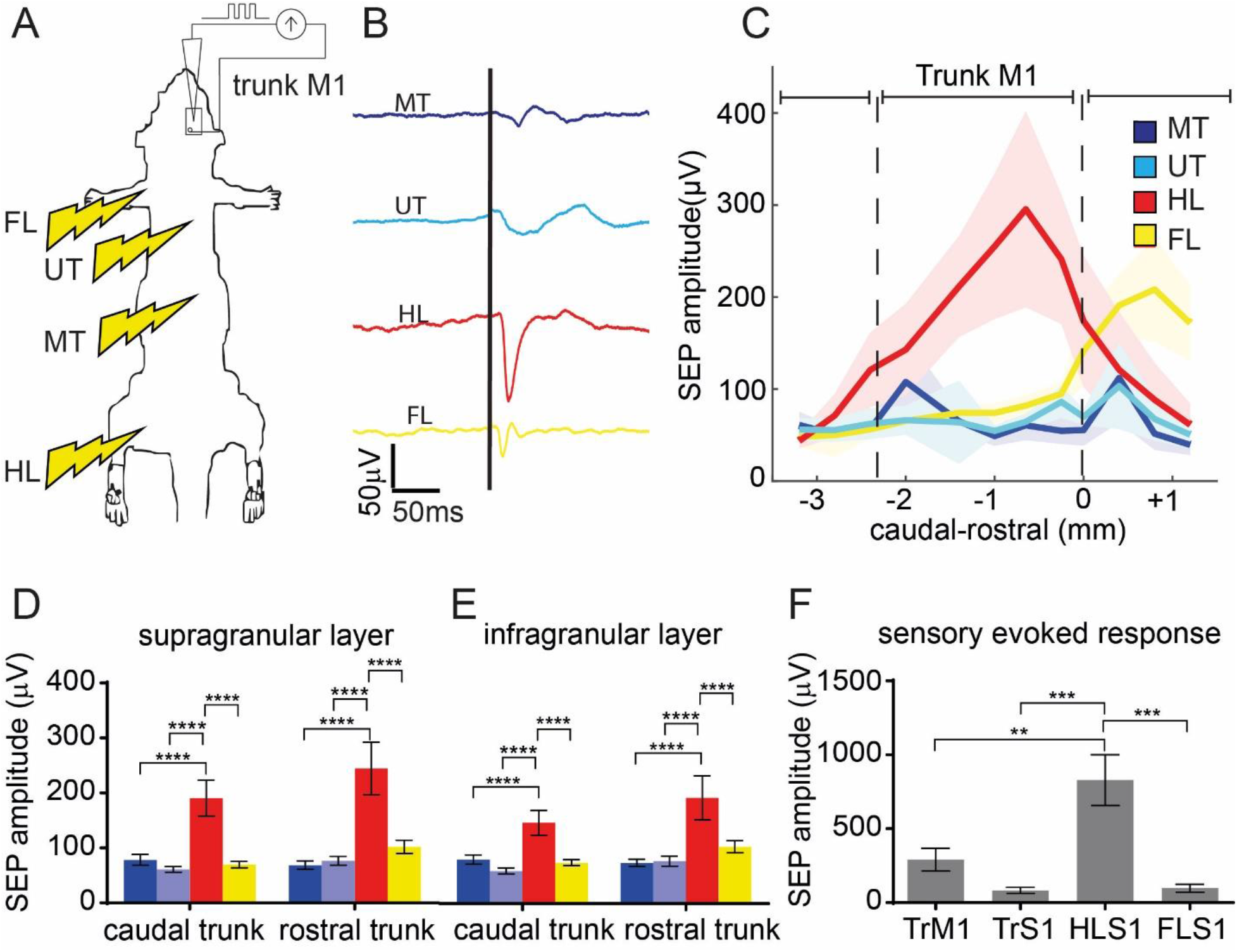
HL proprioceptive information predominates in trunk M1. A) Methodological diagram of the electrical stimulation paradigm. Stimulations occurred in the dorsal hairy skin of forelimb (FL), hindlimb (HL), T4-T5 dermatome (UT), T9-T10 dermatome (MT). B) Example of sensory evoked responses in trunk M1 from the different stimulation locations on the body. C) Sensory evoked potential (SEP) amplitude in the supragranular layer in response to high intensity (5.0mA) stimulation across the rostrocaudal axis of trunk M1 at 1.25 mm lateral from midline. D-E) SEP amplitude in the supragranular (D) and infragranular (E) layers in the caudal region (−1 mm to −2 mm RC relative to bregma) and in the rostral region (−0.75 mm to 0 mm RC relative to bregma) of the trunk M1. F) Sensory evoked response in the supragranular layer to high amplitude (5.0 mA) hindlimb stimulation in the sensory and motor cortices of trunk, hindlimb, and forelimb.

Since there was an internal motor somatotopy along the rostrocaudal axis of trunk M1 (refer to Figure 5F), cortical locations where SEPs were recorded were segregated into rostral (0 to –0.75 mm caudal to bregma) and caudal regions (−1 to −2 mm caudal to bregma). In the supragranular layer, there was no effect of recording location (two-way ANOVA, *F* (1,191) = 2.34, *p* = .13) but there was an effect of stimulus location (two-way ANOVA, *F* (3,191) = 22.25, *p* < .0001; Figure 6D) such that the SEP amplitude recorded from both rostral and caudal trunk in response to HL stimulation was greater than the response to stimulation of all the other stimulus locations (multiple comparison test, *p* < .0001). This result demonstrates an important role for HL sensory integration within trunk M1 but without any somatotopic organization within trunk M1. Surprisingly, there was no difference in the SEP amplitude in response to UT stimulation compared to MT stimulation (multiple comparisons test, *p* = .99) suggesting no somatotopy of trunk sensory input within trunk motor cortex.

In the infragranular layer, there was an overall effect of stimulus location (two-way ANOVA, *F* (3,172) =14.48, *p* < .0001; Figure 6E), where the SEP amplitude to HL stimulation was again greater in both the caudal and rostral region of trunk M1. Moreover, similar to the supragranular layer, there were no differences between SEP amplitude in response to MT stimulation compared to UT stimulation (multiple comparisons test, *p* = .95) suggesting similar organization for both supra and infragranular layers. Finally, to assess the effectiveness of high intensity (5.0 mA) HL sensory stimulation to reach the sensory or motor cortex, the SEP amplitude was compared across trunk M1, trunk S1, hindlimb S1 and forelimb S1. There was an overall effect of cortical location (one-way ANOVA, *F* (3, 21) = 10.14, *p* < .001; Figure 6F) and, as expected, the SEP response in the hindlimb S1 was greater than the response in any other region (Tukey’s multiple comparison, HL S1 vs trunk S1: *p* < 0.001, HL S1 vs trunk M1: *p* < 0.01, HL S1 vs FL S1: *p* < 0.001). These results further support the extensive and preferential integration of hindlimb sensory input into trunk M1.

### Sensorimotor integration is cortico-cortical for trunk stimuli, thalamo-cortical for hindlimb stimuli

Since sensorimotor integration in the cortex is known to be primarily mediated by projections from the S1 cortex and the thalamus (1, 49, 50), retrograde tracing was used to better understand the relative contribution of cortico-cortical versus thalamo-cortical connections to trunk M1 (Figure 7A). Tracing revealed that trunk M1 received cortico-cortical input from ipsilateral trunk S1, hindlimb S1, and forelimb S1. However, the relative contribution from these sensory cortices was highly variable across animals (Figure 7B). For example, rat 1 appeared to have more cells projecting from hindlimb S1 to trunk M1 than from trunk S1 to trunk M1, while rat 3 exclusively showed projections from dorsal trunk S1 to trunk M1. Trunk M1 also received input from secondary sensory cortex, dysgranular zone, shoulder S1, whisker and face S1 (data not shown), thereby making trunk M1 a crossroad for somatosensory information. In all animals, the projections from S1 to trunk M1 were predominantly mediated by S1 cells in the supragranular and infragranular layers (Figure 7C). This laminar specificity is consistent with studies in the whisker sensorimotor system (1, 5). Tracing also revealed strong thalamo-cortical projections, likely carrying proprioceptive information (51), from the ventral posterolateral nucleus of thalamus (VPL) to trunk M1 in all animals (Figure 7D).

**Figure 7.**
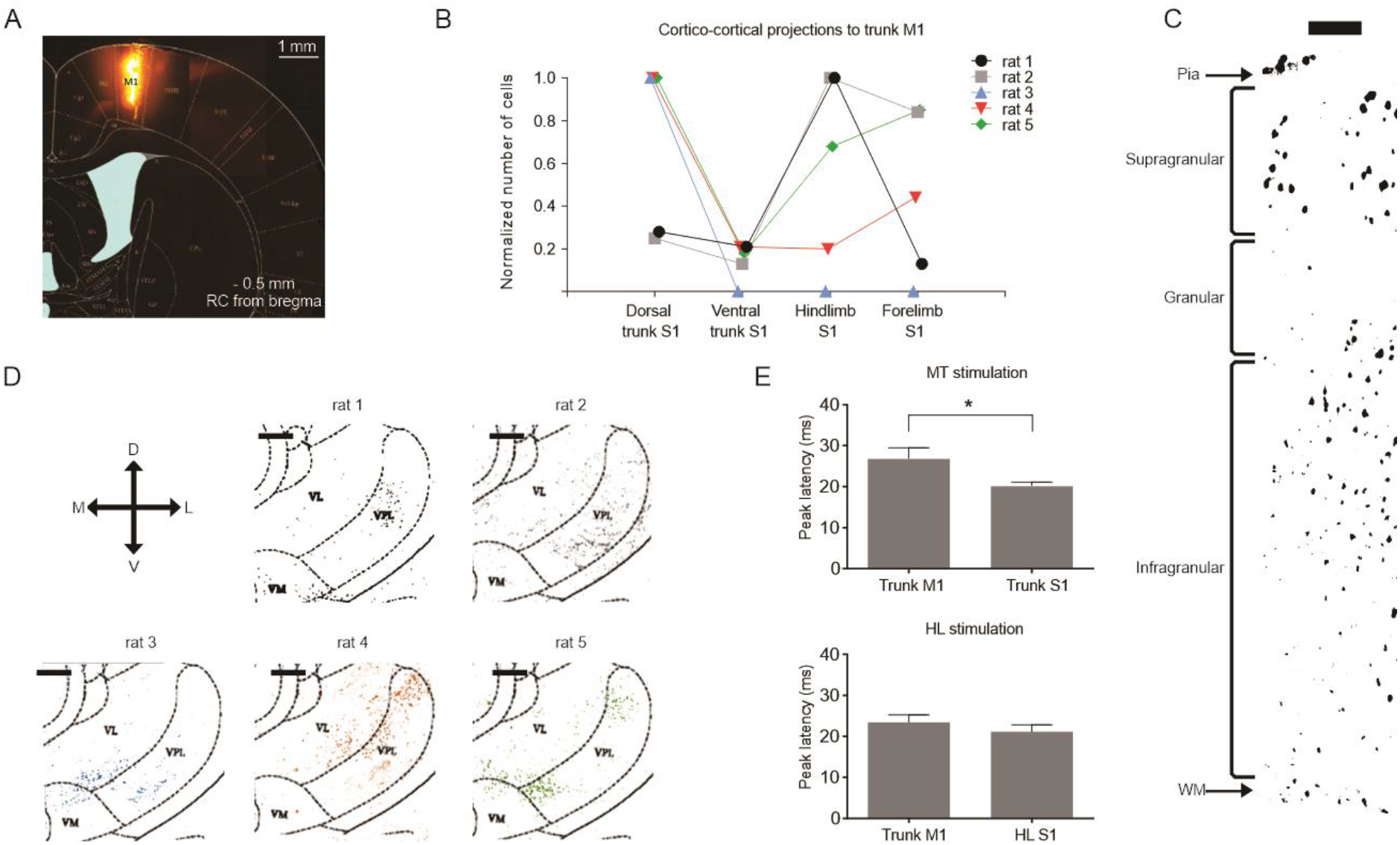
Proprioceptive information from hindimbs arrives in trunk M1 at the same time as hindlimb sensory information. A) Coronal brain slice with superimposed rat brain atlas image (113). The injection site (−0.5 mm RC, 1.25 mm ML, 1.65 mm DV relative to bregma) is limited to trunk M1. B) Normalized number of labeled cells in different sensory cortices (dorsal trunk S1, ventral trunk S1, hindlimb S1 and forelimb S1, for coordinates, see Method). Most of the trunk M1 projecting cells are located in trunk S1 and hindlimb S1. C) Black and white image of the labelled cortical cells in a coronal view of trunk S1. Most of the neurons are located in the supra and infragranular layers. Scale bar: 0.4 mm D) Image of the labelled thalamic cells in a coronal view with the superimposed 2007 Paxinos and Watson atlas. Thalamic neurons are located in the ventro-postero-lateral nucleus (VPL) of the thalamus. Scale bar: 0.2 mm. E) Top: peak latency of the high intensity mid-trunk stimulation in the trunk M1 and trunk S1. Bottom: peak latency of the high intensity mid-trunk stimulation in the trunk M1 and hindlimb S1.

To determine if the source of projections to trunk M1 differed between body parts that were stimulated, SEP latency was analyzed. The latency of the SEP recorded from trunk M1 was significantly later than that of trunk S1 when MT was stimulated. The median latency difference was 5 ms (Mann-Whitney test, *p* < .05; Figure 7E). This led us to conclude that the sensorimotor integration of trunk sensory information in trunk M1 is primarily mediated by cortico-cortical projections. In contrast, the latency of the SEP recorded from trunk M1 and hindlimb S1 in response to hindlimb stimulation was similar (median latency: 21.65 ms) (Mann-Whitney test, *p* = .39; Figure 7E). This led us to conclude that the integration of hindlimb sensory input in trunk M1 is primarily mediated by thalamo-cortical projections, specifically the VPL carrying proprioceptive information. To identify how this sensory information might be used, next we recorded single neurons from trunk S1 and M1 while animals were subjected to tilts in the lateral plane (see next).

### Postural control is predominately supported by hindlimb sensory and lower trunk motor cortices

To investigate the importance of sensorimotor integration between trunk and hindlimb in postural control, we used our tilt task in which animals are subjected to unexpected tilts in the lateral plane (52) while single units were recorded from different primary sensory (forelimb, hindlimb and trunk) and primary motor (hindlimb, trunk) cortices (Figure 8A, 8B). Three measures from the neuronal data were compared: responsiveness (proportion of neurons responding), magnitude of the single neuron response and mutual information carried by the neuronal response regarding the severity of the tilt (Fig. 8C-G). Concerning sensory cortices, trunk S1 was less involved in postural control than either hindlimb or forelimb S1. First, while trunk S1 cells were responsive to the task, both hindlimb and forelimb S1 cells were more likely to respond than trunk S1 cells (HL: 82%, FL: 57%, Trunk: 32%, Chi-square test between HL and Trunk: *p* < .0001, FL and Trunk: *p* < .001, between HL and FL: *p* < .05, Figure 8C). Second, for trunk S1 cells that responded to the tilt, the magnitude of the response was smaller than that of hindlimb or forelimb S1 cells (magnitude of the response: hindlimb S1: 3.32 +/− 0.36 Hz, forelimb S1: 2.94 +/− 0.32 Hz, trunk S1: 1.87 +/− 0.19, Kruskal-Wallis test, *p* < .0001, Dunn’s multi comparison test: hindlimb S1 vs trunk S1 (p < .001), forelimb S1 vs trunk S1 (*p* < .01), Figure 8D). Lastly, trunk S1 neurons were less discriminative of the type of tilt, conveying less mutual information about the severity of the tilt (hindlimb S1: 0.101 +/− 0.020 bits, forelimb S1: 0.086 +/− 0.015 bits, trunk S1: 0.060 +/− 0.007 bits, Kruskal-Wallis test, *p* < .0001, Dunn’s multiple comparison test: hindlimb S1 vs trunk S1 (*p* < .001), forelimb S1 vs trunk S1 (*p* < .01), Figure 8E). Importantly, after dividing trunk S1 into LT, MT, UT (see material and method), there were no differences between these trunk subregions for any measure (responsiveness: Chi-square: *p* = .06, magnitude of response: Kruskal-Wallis test: *p* = .2412, mutual information: Kruskal-Wallis test: *p* = .2925; Figure 8C-E). Therefore, as the animal worked to maintain its balance in response to unexpected tilts, the hindlimb and forelimb sensory cortices provide more information about the tilt than the trunk sensory cortex, likely due to the fact that they are conveying both tactile and proprioceptive input while the trunk is only conveying proprioceptive input.

**Figure 8.**
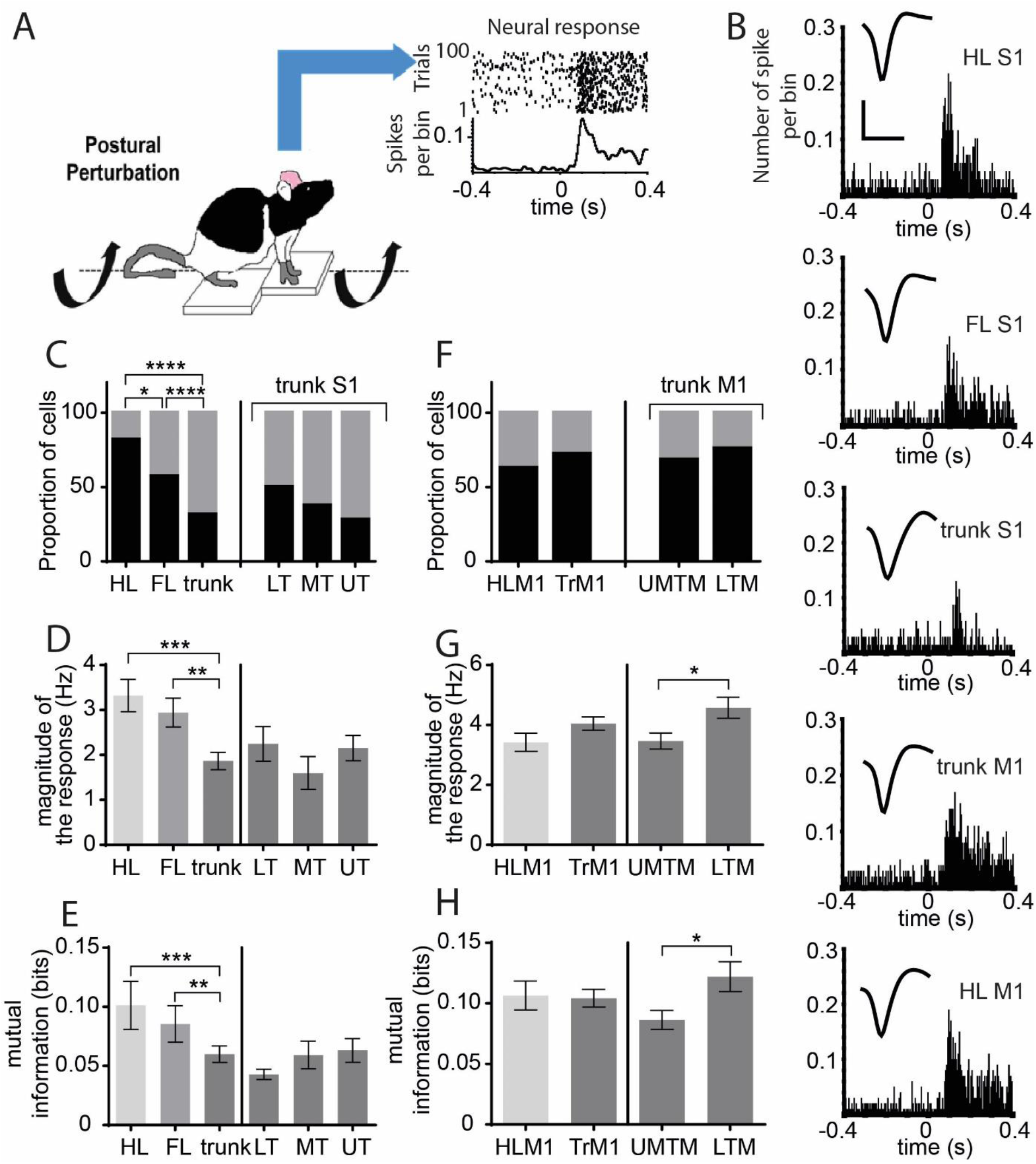
Hindlimb S1 and lower trunk M1 combine to carry the most information about postural control. A) Methodological diagram of the postural control task. The animal is experiencing unexpected tilts in the horizontal plane while single neurons in different sensory and motor cortices are recorded. B) Example PSTHs showing a neuron response to the unexpected tilt for each recorded cortical area in sensory and motor cortices. The waveform scale on top: y-axis: 0.05mV, x-axis: 0.6ms. C) Responsiveness in different sensory cortices. Data presented as: cortical area: (number of responsive cells, number of non-responsive cells, % of responsive cells). Hindlimb S1: (32, 7, 82%), forelimb S1: (39, 29, 57%), trunk S1: (75, 162, 32%), LT: (13, 13, 50%), MT: (31, 51, 38%), UT: (36, 93, 28%). D) Magnitude of the response (for responsive cells only) in different sensory cortices. E) Mutual information in different sensory cortices. F) Responsiveness in different motor cortices. Data presented as: cortical area: (number of responsive cells, number of non-responsive cells, % of responsive cells). G) Magnitude of the response (for responsive cells only) in different motor cortices. H) Mutual information in different motor cortices.

Trunk M1 cells had a tendency to be more responsive to the task compared to hindlimb M1 (responsiveness: hindlimb M1: 63%, trunk M1: 72%, Chi-square test: *p* = .0658, Figure 8F). Moreover, the magnitude of the trunk M1 response was larger than that of hindlimb M1 (magnitude of the response: hindlimb M1: 3.42 +/− 0.30 Hz, trunk M1: 4.05 +/− 0.23 Hz, Mann-Whitney test: p < .05, Figure 8G). Surprisingly, despite these differences in responsiveness, hindlimb M1 and trunk M1 provided similar information about the severity of the tilt (mutual information: hindlimb M1: 0.107 +/− 0.012 bits, trunk M1: 0.104 +/− 0.007 bits, Mann-Whitney test: p = .3987, Fig. 8H). Interestingly, when examining the response of different subregions within trunk M1, lower trunk M1 (LTM) was more involved than upper/mid trunk M1 (UMTM).

In fact, even though neurons in LTM had a similar proportion of cells responding to the tilts compared to UMTM (LTM: 76%, hindlimb M1: 69%, Chi-square test: *p* = .3141; Figure 8F), the magnitude of the response of LTM neurons was greater than neurons in UMTM (LTM: 4.57 +/− 0.35, UMTM: 3.46 +/− 0.27, Mann-Whitney test: *p* < .05; Figure 8G). This resulted in more information about the severity of the tilt being encoded by LTM compared to UMTM (LTM: 0.122 +/− 0.012 bits, UMTM: 0.086 +/− 0.007 bits, Mann-Whitney test: *p* < .05; Figure 8H). These data suggest that the region that activates lower trunk musculature (LTM) may be specialized for postural control.

## Discussion

Together, these data present an extensive view describing how cortical organization is relevant to function by demonstrating how the integration of proprioceptive input from the hindlimbs and trunk within lower trunk M1 can be used for not only locomotion but for a range of functions including postural control. Summarizing, the trunk sensory and motor cortices are larger than previously reported and there is relevant somatotopy within both the trunk sensory and motor cortex. Separately, trunk motor cortex is well integrated with hindlimb motor cortex and trunk sensory information is well integrated into hindlimb sensory cortex, but only for high intensity stimuli that activates proprioceptive pathways. Furthermore, the functional role of this integration of hindlimb and trunk proprioceptive information into trunk motor cortex for postural control was demonstrated by the cortical response to tilts in the lateral plane recorded from awake animals. This has important implication for recovery of function after neurological injury or disease (35, 52, 53) discussed below.

### Methodological Considerations

Choices made in our experimental design impacted data analysis. First, for sensory maps, we chose to record from as many single units as possible, identifying the extent of each cells receptive field as our recording electrode was passed through the entire depth of the S1 cortex. Therefore, it was not possible to sample the entire trunk S1 cortex within a single animal due to time constraints. Similarly, for the trunk M1 cortex, we chose to sample from as many muscles as possible, adding to the length of the surgery, limiting our ability to sample the entire trunk M1 within every animal. Moreover, here we show that, unlike the whisker, forelimb, and hindlimb sensory systems that tend to have differences in RF size across layers (11), the RF size of neurons in trunk S1 was similar across layers, within the same RF center, but the role of urethane anesthesia in this assessment cannot be ruled out (54). Furthermore, we conclude that the response to high intensity stimulation reflected proprioceptive information, while that of low intensity stimulation reflected tactile stimulation. There are three pieces of evidence for this. First, high intensity stimulation generated twitches in the underlying muscles that would activate proprioceptive nerve endings. Second, our comparison of the latency of response and, third, the identification of trunk M1 afferents originating in VPL thalamus, further support activation of proprioceptive information. Therefore, the logical conclusion is that the responses to high amplitude stimulation are due to activation of proprioceptive information, although we did not show this definitively.

### Oppositional gradient in overlap across thoracic dermatomes from DRG to trunk S1

Sensory information from the trunk is differentially integrated at multiple levels along the entire neural axis. At the spinal level, overlap between thoracic dermatomes is graded such that caudal DRGs (T10-T13) have less overlap than rostral DRGs (T1-T5). The functional implication of this is that dorsal rhizotomy of a caudal DRG would result in a more complete deafferentation than a dorsal rhizotomy of a rostral DRG. At the same time, representation of these dermatomes in S1 have the opposite gradient regarding overlap. The RF size of trunk S1 neurons increased along the mediolateral axis of cortex, such that the lower thoracic trunk S1 neurons had a greater RF size compared to the upper thoracic trunk S1 neurons. This change in RF size across trunk cortex was also true for ventral trunk representation (21). Taken together, for a given stimulus to lower trunk dermatomes, the limited overlap produces fewer DRGs conveying sensory information to the cortex where S1 neurons have larger receptive fields, amplifying the signal. In contrast, for upper trunk dermatomes, the greater overlap across DRGs amplifies information to the cortex where neurons have smaller receptive fields. Therefore, the lack of overlap at the spinal level is compensated for by the greater overlap at the cortical layer and vice versa. However, this does not prevent damage at the lower thoracic level from resulting in a greater loss of information than damage at the upper thoracic level. It may be that due to the dexterous use of the forelimbs it is considered more important to preserve upper trunk sensory information than lower trunk.

### Integration of sensorimotor information within trunk M1 is not strictly homotopic

The idea that sensorimotor integration solely takes place in homotopic areas (e.g. whisker M1 would exclusively receive information from whisker S1) has been contested by recent studies. Classical studies showed that neurons in whisker M1, forelimb M1, and hindlimb M1 received sensory input from the same body part that induced movement when activated with ICMS (55–57). However, more recent studies have shown that sensorimotor integration occurs between sensory and motor systems of different body parts that must communicate with each other to produce an optimal behavior. Studies examining cortical control of jaw movement revealed the integration of orofacial sensory input, as an essential component (58–60). Recent evidence also showed that acute proprioceptive stimulation provided to the tooth pulp located in the jaw affects sensorimotor integration in the face motor cortex (61). Taken together, these studies suggest that some sensorimotor systems are integrated on a functional basis more so than an anatomical basis. To our knowledge, our study is the first to bring insight on sensorimotor integration in the trunk motor cortex. This is of fundamental importance since many studies have demonstrated that sensory feedback to the motor cortex is critical during locomotion and recovery of function after spinal cord injury (2, 35, 53, 62–64)

Indeed, low amplitude tactile stimulation of dorsal trunk did not impact trunk M1. On the other hand, high amplitude stimulation of both the forelimbs and hindlimbs, conveying proprioceptive information, did reach trunk M1. In fact, we found that the majority of trunk M1 afferents originated in trunk and hindlimb S1 neurons, as well as the aforementioned VPL thalamus. This preferential integration of hindlimb proprioceptive input to trunk M1 combined with the extensive overlap between hindlimb and trunk M1 demonstrate that sensorimotor integration is not exclusively reserved for corresponding sensory and motor cortices but is rather a process that supports communication between the broader sensorimotor cortex to achieve optimal behavior, as outlined next.

### Trunk sensorimotor integration extends our understanding of trunk M1 as a biomechanical link between forelimb and hindlimb

The present data extend the theory that trunk muscles serve as a biomechanical link between the forelimbs and hindlimbs (65), providing important insight into how this occurs. First, although trunk M1 is defined as the region that most likely activates trunk muscles, this region rarely activated trunk muscles exclusively, but rather is coactivated with hindlimb muscles, and in a more limited fashion, with forelimb muscles. Consistent with previous studies of other rodent species (66, 67), the location of trunk M1 was located medial to both forelimb and hindlimb M1. The area devoted to exclusive activation of trunk was small and its location was inconsistent across animals. This is in agreement with previous literature (26, 31, 32, 34, 35, 46) and is likely due to the fact that these animals were cage-raised (68).

It makes sense that trunk muscles were rarely activated in isolation since, functionally, trunk movement alone is not useful in most behavioral situations but rather the trunk moves in concert with the limbs. The integration supports the role of lower thoracic trunk muscles synergistically acting with the hindlimbs to aid in postural control during locomotion (69). Furthermore, approximately half of the animals had coactivation of forelimb and hindlimb without concomitant activation of trunk muscles. The area was located within the synergistic trunk region (data not shown). This movement representation was also found in other species across phylogenic scales such as mouse (70), tree squirrel (71), tree shrew (66), prosimian Galagos (72) and macaque monkey (73). These synchronous forelimb-hindlimb coactivations (74) might be part of a range of movement types associated with locomotion, including for example galloping (75), that require a linkage between forelimb and hindlimb.

Finally, sensorimotor integration is required for proper functioning. For example, here we show that high intensity midtrunk stimulation activating proprioceptive sensors resulted in greater activation of hindlimb S1 than forelimb S1. This preferential integration between hindlimb S1 and trunk proprioceptive information indicates that hindlimb S1 is modulated by the location and movement of trunk in space. This proprioceptive information would thus guide the lower limbs during locomotion (2).

### Trunk sensorimotor integration supports a range of functions

Trunk M1 not only modulates limb movement, but our data supports the known role of trunk M1 in other functions. Trunk M1 is known to be involved in the control of autonomic function. In fact, it has been shown that the trunk cortical representation overlaps with parts of neocortex that indirectly influence visceromotor function (76). Moreover, ventral trunk has a role in controlling muscles that are involved in bladder pressure control (77) and voiding behavior (78). The extensive integration of hindlimb sensory information with trunk M1 is likely used to support these functions.

Moreover, while the rostral portion of trunk M1 activated upper thoracic trunk muscles, the caudal portion was involved in controlling lower thoracic trunk musculature. It has been previously shown that these lower thoracic trunk muscles play an important role in sexual posturing and lordosis as observed in the female rat (79). Our results showed that the caudal portion of trunk M1 that controls these lower thoracic muscles overlaps with the genital motor cortex (23). Moreover, we show here that the entire extent of trunk M1 receives proprioceptive trunk information from upper and mid trunk, demonstrating extensive integration of trunk proprioceptive information across the trunk motor cortex that could be useful for sexual posturing.

### Role of trunk sensorimotor cortex in postural control

The hindlimb-trunk M1 coactivation zone identified in this study extends our previously described role of hindlimb M1 cortex in postural control (52, 80). For the sensory cortex, previous studies showed that tactile and proprioceptive somatosensory feedback from the peripheral limbs are involved in postural control (81, 82). This is consistent with our data here showing that for both hindlimb and forelimb S1, more cells respond and the magnitude of the neural response is greater compared to that of trunk S1, such that forelimb and hindlimb trunk S1 convey more information about the tilt than that of trunk S1. But, within trunk S1, the different areas of trunk (LT, MT and UT S1) are equally responsive, conveying similar amounts of information about tilt. These difference between forelimb and hindlimb S1 on the one hand and trunk on the other, is likely mediated solely by proprioceptive information with little tactile information since the trunk is not in contact with the platform. Indeed, while the limbs of the animal are in contact with the moving platform providing ascending tactile information this is not true for the trunk. Furthermore, since, in this study, we showed that a significant proportion of neurons in trunk S1 responded to hindlimb proprioceptive information, during the tilt task, the proprioceptive information reaching trunk S1 comes from the position of both the trunk and hindlimb in space, allowing significant integration of this proprioceptive information to allow the animal to maintain its balance.

In the motor cortex, a similar proportion of hindlimb and trunk M1 cells were likely to respond with a similar magnitude of response, conveying a similar amount of information about the tilt, suggesting that these two regions are equally active in controlling muscles when the animal is maintaining its balance in response to the tilt. Interestingly, within trunk M1, the region that controls lower thoracic muscles (caudal trunk M1) was more engaged in the task compared to the regions that control upper and mid thoracic musculature (rostral trunk M1). Given that more of the weight of the animal is over the hindlimbs, these data suggest that extensive integration across hindlimb and lower thoracic trunk M1 is used for postural control.

Our results also support a specialized and sensitive role for lower thoracic trunk M1 to detect changes in posture driven by movement of the pelvis in space in coordination with hindlimbs. This is consistent with the discussion of sexual posturing, discussed above and is supported by our mapping studies that showed proprioceptive information from hindlimbs and trunk arrives simultaneously in lower trunk M1 directly via thalamus. These data clarify the importance of sensorimotor integration between hindlimb and the lower body for postural control.

### Hindlimb sensory feedback to the trunk sensorimotor cortex: Pathophysiological implications

Pathologies resulting in postural deficits in humans are associated with changes in cortical organization(83), motor planning (84) and recruitment of trunk musculature (85). The integration of hindlimb proprioceptive information in trunk M1 cortex elicited here provides an opportunity for a new understanding of how therapy after mid thoracic spinal cord injury improves function. For a complete spinal transection, we previously showed that therapy produced sprouting of descending corticospinal axons from hindlimb M1 cortex into thoracic spinal cord that could be used to control trunk musculature. This produced a larger representation of the trunk M1 cortex whose extent was correlated to functional recovery and overlapped with expansion of the forelimb sensory cortex, creating a new circuit of forelimb sensory and trunk motor integration (34). If this reorganized cortex was lesioned, functional gains were lost (35). Our new understanding of the extensive sensorimotor integration in intact animals presented here makes it more clear that the sensorimotor integration in animals that receive therapy after SCI is not a novel sensorimotor integration but a necessary restoration of a system that operates on strong sensorimotor organization.

While the role of limb proprioception after more severe injuries is less understood, our group previously showed that after complete spinal transection, when sensory input from the hindlimb is not possible, epidural stimulation induces sensory feedback into the deafferented hindlimb sensorimotor cortex that carries information about the animal’s behavior (53). This study now makes clear that this sensory feedback is likely to be trunk proprioceptive information that provides input to hindlimb M1 cortex in intact animals. Therefore, therapy to improve function could take advantage of this pre-existing sensorimotor integration to restore function.

This role of sensorimotor integration extends to models of partial spinal lesion. Proprioceptive information has been suggested to be critical for recovery of function after mid thoracic spinal cord injury (86). Follow-on epidural stimulation of spinal circuitry below the level of the lesion restored volitional locomotion in rats(53, 87, 88), non-human primates (89) and humans (90–92). Stimulation is conducted at lateral sites, over or near the DRGs and it has been suggested that this epidural stimulation activates proprioceptive afferents (92, 93).

At the same time, anatomical studies have suggested that this proprioceptive input from below the level of the lesion interacts with extensive reorganization of the spinal proprioceptive circuitry (93–96) that can allow the proprioceptive input to reach the cortex through interactions with reorganized supraspinal circuits (87, 97–100). Therefore, the hindlimb proprioceptive information that is enabled through epidural stimulation replaces the hindlimb proprioceptive information we now know to exist in intact animals.

Therefore, the work outlined in this paper supports the idea that sensorimotor integration across broad regions of the cortex is key to improving treatment outcomes after neurological damage or disease (101) and now we understand that this sensorimotor integration is the operational model of the trunk cortex in intact animals. Moving forward, our understanding of the sensorimotor integration in the intact system could be used to tailor rehabilitative strategies to optimize sensorimotor integration or functional recovery.

## Material and Methods

### Subjects

One hundred and six adult, female Sprague Dawley rats (225-250 g; Envigo) were maintained on a 12/12-hour light/dark cycle with ad libitum food and water. Fifteen animals were used to map the representation of each thoracic dermatome at the spinal level, 40 animals were used to map the internal representation of the trunk sensory cortex, 21 animals were used to examine the movement representation of trunk motor cortex, 14 animals were used to examine the integration of sensory information within and between sensory and motor cortices, five animals were used for anatomical tracing, and 11 animals were used to study sensorimotor integration relevant for postural control.

For all anesthetized experiments, animals were secured on a stereotaxic frame (Neurostar, Sindelfingen, Germany) and body temperature was maintained at 37°C using a temperature-controlled heating pad (FHC Inc., Bowdoin, ME). In addition, heart rate, SpO2, and anesthetic state (whisking/toe pinch reflex/corneal reflex) were constantly monitored. All experimental procedures were approved by UC Davis or Drexel University IACUCs and followed NIH guidelines.

### Body grid system to map receptive fields

To identify receptive fields (RFs) consistently across animals, the dorsal trunk was shaved and a grid of 128 equally spaced squares (8 columns and 16 rows) was drawn indelibly. The grid spanned from the skull’s base, parallel to the intertragic notch of the ear, to the tail’s base, and from the dorsal trunk’s midline to its lateral aspect parallel to the knee (Figure 1A). Each grid square was approximately 1 cm^2^ and was consistent across animals due to the similarity of both size and weight. In addition, a photograph of the animal with the drawn grid was taken in order to assist in defining RFs during sensory mapping experiments.

### Mapping thoracic dermatomes

Animals were anesthetized with urethane (1.5 g/kg, IP) and maintained at Stage III-3 anesthesia (54). An incision was made along the midline of the trunk, and axial musculature was separated from the vertebral column to expose the thoracic vertebrae. The spinous processes, lamina, and transverse processes of the selected thoracic vertebrae were carefully removed to access the dorsal root ganglion (DRG) on one side of the body. The animal’s spinal column was secured in place by attaching locking forceps to the transverse process rostral to the T1 vertebrae and caudal to the T13 vertebrae. A single high-impedance (4-10 MΩ) tungsten microelectrode (FHC Inc., Bowdoin, ME) was attached to the stereotaxic manipulator and a ground wire was placed in contact with the body cavity. The electrode was positioned over a single DRG and lowered slowly until a single cell was identified. The neuronal signal (digitized at 40 kHz) was amplified (20000x), band pass filtered (150 - 8000 Hz, Plexon Inc., Dallas, TX) and monitored with an oscilloscope and through audio speakers. The cell’s receptive field was then identified using light tactile stimulation (11, 19). First, the dorsal cutaneous surface of the animal was tapped with a cotton brush, both within and outside the body grid to gain insight of the neuron’s RF. If the RF was located within the trunk body grid, it was then mapped with a 0.25 body grid square resolution by applying light tactile stimulation to the cutaneous surface using a wooden probe (4 mm diameter). If the RF was found outside the grid, it was not included in any further analysis. When mapping of that neuron’s RF was complete, the electrode was lowered at least 50 μm dorso-ventral (DV) before another cell was identified to ensure that the same cell was not mapped twice. This process was repeated until the electrode punctured through the entire DRG. Each DRG was sampled at least three times, so as to cover the DRG rostrocaudal extent (38). A trunk dermatome was defined as the union of all trunk grid locations on the skin that were found to be responsive to at least one cell in the respective DRG. The width of a dermatome was defined as the number of trunk grid locations within its rostrocaudal extent. Center position of a dermatome was defined as the center of this rostrocaudal extent. Dermatomal overlap was defined between two adjacent dermatomes as the distance between the rostral extent of the more caudal dermatome and the caudal extent of the more rostral dermatome.

### Mapping trunk sensory cortex (S1)

Animals were anesthetized with urethane (1.5 g/kg, IP) and maintained at Stage III-3 anesthesia (54). A craniotomy was performed on the right hemisphere to expose the hindlimb, trunk and parts of the forelimb primary sensory cortices (S1) (19, 102). Based on a pilot study (*n*=3), 80 predefined cortical locations were chosen. They extended from −2.0 mm to −3.8 mm rostrocaudal (RC) from bregma with a resolution of 0.2 mm, and from −2.0 mm to −3.75 mm mediolateral (ML) with a resolution of 0.25 mm between locations. At each location, the electrode waslowered into the brain up to a depth of −2.0 mm dorsoventral (DV) while light tactile stimulation was applied. If a neuron was responsive, the neuron’s receptive field was categorized into either trunk, ventral trunk, head/face, forelimb, hindlimb, tail, or a combination of body categories. If the RF included the trunk, the RF was further analyzed relative to body grid with a 1.0 body grid square resolution and calculated separately for the supragranular, granular, and infragranular layers. A somatotopic map of the trunk and surrounding sensory cortices was constructed. At each cortical location, the proportion of cells responding to each body category was investigated. A body category was assigned to a cortical location if at least 25% of the neurons in that location were responsive to that body category. If there were multiple body categories that meet the criterion, the category with the highest proportion of responsive neurons was chosen (Fig. 2B).

In order to locate the cortical representation of the thoracic dermatomes within the trunk S1, all cells that had RF centers within trunk were used. For a given cortical location, the rostrocaudal positions of the RF centers on the body grid from all cells of that location were averaged. The dermatome with the closest center position to the average cortical RF position defined the corresponding dermatome of that cortical location. Cortical locations that represented the same dermatome were grouped to generate the representation of thoracic dermatomes in the cortex. All neuron’s RF belonging to the same dermatome representation were used to calculate the amount of overlap between the neighboring dermatome representations‥ To analyze size and extent of trunk RFs, only neurons that were completely contained within the borders of the trunk grid were used. Average RF size was calculated by averaging the number of responsive body grid squares for all cells.

### Local field potential recording in response to peripheral electrical stimulation

Animals were anesthetized with urethane (1.5 g/kg, IP). A craniotomy was performed on the right hemisphere to expose the sensory and motor cortices. A 32-channel, four shank recording electrode array (A4×8-5mm-200-400-177, NeuroNexus, Ann Arbor, MI) was positioned over the fixed locations either spanning the trunk M1 (−3.2 to 1.2 mm RC, 1.25 mm ML), trunk S1 (−3.4 mm to −2.2 mm RC, 3 mm ML), hindlimb S1 (−1 mm to −2.2 mm RC, 2.5 mm ML), or forelimb S1 (+0.5 mm to −0.7 mm RC, 3.5 mm ML). The array was lowered perpendicularly into the cortex to a depth of 1.8mm where it was fixed in place. Bipolar stimulating electrodes were inserted subcutaneously into the dorsal hairy skin at four locations: hindlimb (HL), wrist of the forelimb (FL), T4-T5 dermatome of the upper trunk (UT), and T9 dermatome of the mid trunk (MT), contralateral to the recording location. Electrical stimulation, consisting of 100 pulses (1ms duration) was delivered every two seconds at varying stimulation intensities (0.5 or 5 mA). A low intensity stimulus (0.5 mA) was used to activate the low threshold primary afferents (tactile), while high intensity stimulation (5.0 mA) was used to elicit muscle twitches and slight movement to activate proprioceptive afferents and nociceptive afferents (103, 104). Therefore, low intensity stimulation (0.5 mA) mimicked light tactile stimulation while high intensity stimulation (5.0 mA) was used to mimic proprioceptive stimulation. The extracellular local field potential (LFP) was acquired simultaneously from all 32 channels (INTAN Technologies, Los Angeles, CA), digitized at 20 kHz, amplified (192x) and band pass filtered (0.1 Hz – 7.5 kHz).To ensure fair comparisons between the stimulation responses of different locations on the body, the responses of each region to stimulation of their RF centers were compared (RF center identified using light tactile stimulation, see above). A high pass filter of 5 Hz was used to mitigate slow wave activity that developed under urethane anesthesia (105, 106) in the cortical local field potential (LFP). A window of 1s centered on the stimulation time was extracted from the high pass filtered LFP data (5 Hz, Butterworth order 2, zero lag) of each recording site. The data in that window was then averaged across stimulation trials to obtain the somatosensory evoked potential (SEP). A representative channel from the supragranular (400 μm DV), granular (800 μm DV), and infragranular (1200 μm DV) cortex was selected for further analysis. For each layer, the SEP was considered responsive if the amplitude exceeded the mean background activity by three standard deviation. SEP amplitude was evaluated as the absolute value of the first negative peak of the SEP, normalized to the background activity. Peak latency of the SEPs was calculated, as the time of the SEP peak amplitude post stimulus. Only responsive SEPs with latency smaller than or equal to 50ms were considered for further analysis to capture the short latency response. In addition, the LFP from each electrode was filtered (300-8000 Hz) and single neurons were discriminated using PCA analysis and visual inspection using offline sorter (Plexon Inc., Dallas, TX).

Single neuron spike times were used to construct peri-stimulus time histograms (PSTH) to determine the magnitude of the response of a neuron to the peripheral electric stimulation using previously published methods (12, 18, 35, 107, 108). The PSTH consisted of spike counts within 5 ms bins averaged across 100 trials within a window of 100 ms from the time of stimulus (Fig. 3E). A neuron was considered responsive if at least two consecutive bins in the PSTH exceeded three standard deviations above the background window. Response magnitude and the proportion of responsive neurons were quantified across all layer in S1.

### Mapping trunk motor cortex (M1)

The representation of the trunk M1 was examined by analyzing evoked movement and EMG activity in response to stimulation of infragranular neurons in M1. Animals were anesthetized with ketamine (63 mg/kg, IP), xylazine (6 mg/kg, IP) and acepromazine (0.05 mg/kg, IP). The animal was maintained at light Stage III-2 anesthesia throughout the entire mapping procedure (54, 109). Rats were administered dexamethasone (5 mg/kg, IM) to control blood pressure and brain swelling. Supplemental doses of ketamine (20 mg/kg) were administered when necessary. Animals were placed in a stereotaxic frame in a prone position such that the limbs could hang freely. Eight bipolar intramuscular electromyogram (EMG) electrodes (stainless steel; 7 strands; AM-Systems Inc.; Sequim, WA) were implanted on dorsal (longissimus) and ventral (external oblique) trunk muscles at the upper thoracic (T4-T5), mid thoracic (T9-T10) and lower thoracic (T12-T13) levels. One EMG electrode was implanted in each of the contralateral shoulder/trunk (spinous trapezius), contralateral forelimb (forelimb bicep), contralateral hindlimb hip (gluteus maximus) and hindlimb ankle (tibialis anterior) (Fig. 4A). Based on previous studies on rats (27, 33, 34), a craniotomy exposed the medial post bregma area and the caudal forelimb area (−3.5 mm to 1 mm RC and 1 mm to 3 mm ML). Similar to the sensory mapping procedure, 88 predefined cortical locations were chosen spanning the craniotomy. A low impedance glass insulated tungsten electrode (100-500 kΩ; FHC Inc., Bowdoin, ME) attached to a stereotaxic manipulator was inserted into one of the 88 predefined cortical locations. In order to assess microstimulation waveform quality, voltage drop across a 10 kΩ resistor interposed in series between animal ground and the isolated current pulse stimulator (A-M systems, Model 2100, Sequim, WA) was monitored with an oscilloscope. At each M1 location, the electrode was lowered to the infragranular layer (1.5 mm DV) and a long train ICMS was applied (110, 111), consisting of 0.2 ms cathodal leading bipolar current pluses (10 to 100 μA) delivered at 333 Hz for 300 ms. The stimulation current was gradually increased in steps of 10 μA until a reliable movement or EMG response was found.

EMG signals and current stimulus times were sent to a data acquisition system (Intan technologies, Los Angeles, CA). EMG was sampled at 5 kHz, zero-lag band pass filtered (40-400 Hz) and rectified. An EMG envelope was obtained by further filtering the data (zero lag Butterworth low pass filter, 20 Hz, 5th order). The EMG envelope was normalized to its peak value to account for changes in EMG response due to electrode placement, impedance mismatch, signal to noise ratio and muscle size (112). Motor evoked potentials (MEPs) were then obtained by averaging the processed EMG over a time window of 1 s centered on the current stimulus timestamps. If the amplitude of a MEP exceeded the background EMG activity by five standard deviations, it was considered a responsive EMG. The minimum current required for eliciting a movement/EMG response was defined as the threshold current for that cortical location. Once a reliable threshold current was found, the current was increased to 100 μA (suprathreshold), and the movement and EMG responses were recorded. A minimum of five separate stimulations were performed in every cortical location. If no movement or EMG response was evoked at suprathreshold current, the cortical location was determined non responsive. If there were more than three consecutive non-responsive locations, the closest responsive location was rechecked to identify the limits of motor cortex. A combination of visual observation of movements and reponsive EMG locations were used to classify cortical locations into movement types (Table 1). Recruitment of trunk musculature via stimulation of the trunk M1 was examined based on EMG response. Trunk musculature responses were classified into different categories based on the location of responsive trunk EMG along the thoracic level at both threshold and suprathreshold currents (Table 2, Figure 5A). At threshold, the proportion of responsive EMG was compared across thoracic levels.

**Table 1:** Movement type classification

| Movement type | Visual Observation | EMG response |
| --- | --- | --- |
| Forelimb (FL) | Isolated movement of forelimb (wrist or multi joint) | Exclusive EMG response- FL muscle (Fl bicep) |
| Hindlimb (HL) | Isolated movement of hindlimb (digits or multi-joint) | Exclusive EMG response- HL muscles (Gluteus, Tibialis) |
| Forelimb Trunk (FLT) | proximal shoulder movement | Coactivation of FL muscle and Trunk /shoulder muscles |
| Exclusive Trunk (ET) | Isolated movement of thoracic girdle | Exclusive EMG response – Trunk muscles |
| Hindlimb Trunk (HLT) | Movement of hindlimb knee /ankle along with thoracic girdle | EMG response -Trunk muscles |
| Synergistic Trunk (FHT) | Forepaw and HL ankle dorsiflexion movements with trunk adduction | EMG response – Trunk muscles |
| Forelimb-Hindlimb (FLHL) | Exclusive Forepaw and HL ankle dorsiflexion movements | Coactivation of forelimb and hindlimb muscles |
| Whisker (WH) | Whisker movements | Absence of any EMG response |

Explanation of how movement types were determined for each intracranial microstimulation trial.

**Table 2:** Trunk musculature classification

| Trunk musculature type | EMG Response |
| --- | --- |
| Dorsal Trunk | Activation of Spinous trap or longissimus muscles at upper, mid thoracic or lower thoracic level |
| Ventral trunk | Activation of External Oblique muscles at upper, mid or Lower thoracic level |
| Upper thoracic trunk (U) | Activation of Spinous trapezius (SP) or Left thoracic longissimus (LMT) or right thoracic longissimus (RMT) or upper external oblique (UOB) |
| Mid thoracic trunk(M) | Activation of left mid thoracic longissimus (LMT) or right mid thoracic longissimus (RMT) or mid thoracic external oblique (MTO) |
| Lower Thoracic trunk(L) | Activation of Lower thoracic oblique (LTO) or lower thoracic longissimus (LT) |

Muscles that were activated in response to intracranial microstimulation were classified as one of five muscle types.

The muscle responses (muscle identification and movement type) associated with the stimulation of each cortical location were used to calculate a responsiveness score (34, 48). For each movement type (or trunk musculature type), the proportion of responses in each location was determined and transformed to a score as follows: range of 0, 1-33%, 34-66%, and 67-100% a score of 0, 1, 2, or 3, respectively. A score of 0 meant that no movements and no EMG response were recorded and a score of 3 meant that the muscle movement (or EMG response) was elicited for 67 −100% of the cortical stimulation. The average responsiveness score for a specific movement type and/or EMG response was calculated by averaging the score across cortical locations. To control for the fact, that not every cortical location was sampled equally, a responsiveness score was included in the analysis only if there were at least five penetrations in a given location.

### Retrograde tracing

Animals were anesthetized with ketamine (63 mg/kg, IP), xylazine (6 mg/kg, IP), and acepromazine (0.05 mg/kg, IP). A craniotomy was made over trunk M1 (−0.5 mm RC, 1.25 mm ML, 1.65 mm DV, relative to bregma) and 300 nL of 10% fluorescent microbeads (Lumafluor Inc., Naples, FL.; Figure 4D, Figure 7A) were injected with a Hamilton syringe (Tip diameter: 0.1 mm). Three days after the injection, animals were perfused with saline followed by 4% PFA and brains were removed. 50μm coronal sections were mounted under Permount (Fischer Chemical, Geel, Belgium) on microscope slides. Brain slices were then imaged using a wide field microscope (5x/.012 numerical aperture; ZEISS, Oberkochen, Germany) and cell counting was performed using ImageJ (National Institutes of Health, Bethesda, MD). Images were transformed to an 8-bit grey scale image and thresholding was done to minimize artifacts caused by auto fluorescence. Automated cell counting (minimum size: 100 pixels^2^) was conducted in the region of interest (ROI). The different ROIs, corresponding to the different sensory cortices were identified based on electrophysiological sensory mapping data (Figure 2B). For locations outside of trunk S1, ROIs for hindlimb and forelimb sensory cortices were identified based on (102). Only ipsilateral projections were identified. The location of thalamic nuclei was identified by superimposing our images on to the rat brain atlas (Paxinos & Watson, 2007).

### Postural control task (Tilt task)

The tilt task was used to understand the sensorimotor integration in the cortex relevant for postural control. Microwire arrays (32 channel each (8*4), 250-micron resolution, Microprobe. Inc) were implanted in the infragranular layer of the cortex bilaterally spanning trunk S1, hindlimb S1, forelimb S1 on the left hemisphere and trunk M1 and hindlimb M1 on the right hemisphere. For chronic microwire implantation refer methods (52, 114, 115). Single neuron activity was recorded from the different cortices, in response to sudden unexpected postural perturbation in the lateral plane with varied tilt severity (direction (clockwise, counterclockwise), speed (slow (12.5°/s), fast (67.9°/s)). The task engaged the cortex bilaterally and was adapted from (52). For each of the electrode spanning the sensory cortex (left hemisphere), the corresponding receptive field center was identified in response to peripheral electric stimulation (0.5mA, tactile) of the different body parts (FL, HL, UT, MT) and LT (T11-T12 dermatome). The electrode was assigned a region based on the receptive field center (highest SEP amplitude). Electrodes spanning caudal trunk M1 (−1 mm to –2 mm RC,1.25 mm to 1.5 mm ML, refer Figure 5F) were in regions that preferentially control thoracic trunk musculature, defined as lower thoracic trunk M1 (LTM). Electrodes rostral to LTM (0 mm RC to −0.75 mm RC, 1.25 mm to 2.0 mm, in addition to controlling lower thoracic trunk muscles, were also likely to control upper, and mid thoracic trunk musculature. This region is defined as UMTM. Regions lateral to LTM (−1 mm to –2 mm RC,1.75 mm to 2 mm ML) preferentially controlled hindlimb musculature, defined as hindlimb M1 (HL M1).

Responsiveness in the different cortices was calculated as the proportion of responsive neurons to at least 1 tilt type. A neuron was considered responsive to a tilt if the neuronal activity in the response window (400ms from start of tilt) was significantly different from the background (t-test) and there are at least 5 consecutive bins (bin size 5ms) in the response window that exceed the background activity by 2 standard deviation. The magnitude of response (hertz) is defined as the change in the average neuronal firing rate from the background (average firing rate in response window – average background firing rate). Shanon’s mutual information was used to quantify the information about the tilt type provided by the neuronal response of each single neuron within the regions (116). If the neuronal response and the tilt type were completely independent from each other, mutual information is 0 bits, if they are perfectly correlated the mutual information is defined by the entropy of stimulus (tilt type) and would contain 2.0 bits (i.e., log_2_(4), n=4 tilt types) of information.

## Acknowledgements

This work was supported by grant R01NS096971 from the National Institutes of Health and grant 1933751 from the National Science Foundation.

---

**Supplemental Figure 1.**
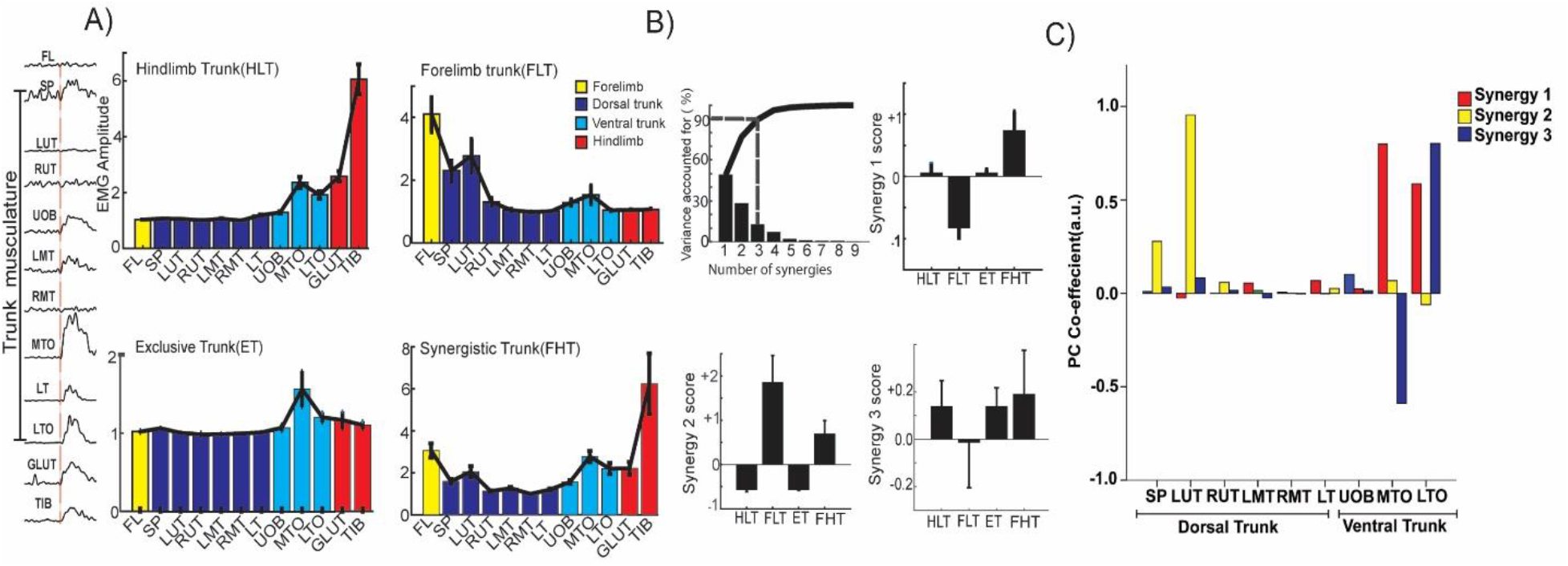
Muscle synergy analysis of the different co-activation zones in trunk M1. A) Average MEP amplitude within the different co-activation zones. EMG electrodes implanted along the trunk, forelimb and hindlimb musculature (Top to bottom: Forelimb bicep (bicep), Spinous trapezius (SP), Left upper thoracic longissimus (LUT), Right upper thoracic longissimus (RUT), Upper external oblique (UOB), Left mid thoracic longissimus (LMT), Right mid thoracic longissimus (RMT), Mid thoracic external oblique (MTO), Lower thoracic longissimus (LT), Lower thoracic external oblique (LTO), Gluteus maximus (Glut), Tibialis anterior (Tib)). B) PCA analysis done to identify the synergies and their synergy scores are plotted. C) Synergy weights: Relative contribution of every trunk muscle towards synergy plotted. Three synergies were extracted that accounted for 90% of the cumulative variance in the data. Synergy 1 and synergy 3 represented ventral trunk musculature activation with the highest weights representing mid and lower thoracic muscles, synergy 2 represented dorsal trunk musculature activation with the highest weights representing the contralateral upper thoracic longissimus. The synergy scores were compared across co-activation zones to explain the role of the different muscle groups. For example, synergy 1 and 2 explain the difference between FLT and HLT co-activation zones (one-way ANOVA F [3,232] = 2.993, p<0.05 for synergy 1 and F [3,232] =14.64, p<0.05 for synergy 2]. Specifically, synergy 1 score of FLT was different from that of HLT (TEST, p<0.05) by almost exclusive contribution of mid and lower thoracic ventral trunk musculature (Supplementary Figure 1). Alternatively, the scores for synergy 2 were different between all co-activation zones (p<0.05), primarily through contribution of upper thoracic dorsal trunk musculature. Synergy 3 scores, whose predominate contribution was also from mid and lower thoracic ventral trunk, were not different between co-activation zones suggesting it might not be possible to explain differences between muscle contribution to exclusive trunk and contribution to co-activation zones.

